# Distinctive and complementary roles of E2F transcription factors during plant replication stress responses

**DOI:** 10.1101/2023.01.06.523015

**Authors:** Maherun Nisa, Thomas Eekhout, Clara Bergis, Jose-Antonio Pedroza-Garcia, Xiaoning He, Christelle Mazubert, Ilse Vercauteren, Rim Brik-Chaouche, Jeannine Drouin-Wahbi, David Latrasse, Catherine Bergounioux, Klaas Vandepoele, Moussa Benhamed, Lieven De Veylder, Cécile Raynaud

## Abstract

Survival of living organisms is fully dependent on their maintenance of genome integrity, being permanently threatened by replication stress in proliferating cells. Although the plant DNA damage response (DDR) regulator SOG1 has been demonstrated to cope with replicative defects, accumulating evidence points to other pathways functioning independently of SOG1. Here, we have studied the role of the Arabidopsis E2FA and EF2B transcription factors, two well-characterized regulators of DNA replication, in the response to replication stress. Through a combination of reverse genetics and chromatin-immunoprecipitation approaches, we show that E2FA and E2FB share many target genes with SOG1, providing evidence for their involvement in the DDR. Analysis of double and triple mutant combinations revealed that E2FB, rather than E2FA, plays the most prominent role in sustaining growth in the presence of replicative defects, either operating antagonistically or synergistically with SOG1. Reversely, SOG1 aids in overcoming the replication defects of E2FA/E2FB-deficient plants. Our data reveal a complex transcriptional network controlling the replication stress response, in which both E2Fs and SOG1 act as key regulatory factors.

**ONE-SENTENCE SUMMARY:** The Arabidopsis E2FA and EF2B transcription factors differently contribute to the plant’s response to DNA replication defects in a cooperative way with the DNA damage response regulator SOG1.

## Introduction

In all eukaryotic organisms, faithful transmission of genetic information from one generation to the next strongly depends on accurate DNA replication. Several factors, including pyrimidine dimers, unrepaired DNA lesions, RNA–DNA hybrids and the formation of DNA secondary structures, can disrupt or slow down DNA replication. These factors can result in fork stalling, leading to replication stress that may in turn affect genomic integrity (Mazouzi et al., 2014). Due to the multiplicity of factors that can lead to fork stalling, replication stress is a ubiquitous threat to the maintenance of genome integrity in all proliferating cells. Interestingly, in plants, there is accumulating evidence that exposure to abiotic or biotic stresses can trigger the DNA damage response (DDR) (Nisa et al., 2019; Pedroza-Garcia et al., 2022), and this effect could partly be due to an increased replication stress (Nisa et al., 2021).

In eukaryotes, the DDR signaling cascade is largely conserved and activates checkpoints that induce cell cycle arrest until the damaged DNA is repaired. The DDR is activated by DNA double-strand breaks (DSBs) or replication stress, and relies on the two protein kinases ATAXIA TELANGIECTASIA MUTATED (ATM) and ATM AND RAD3-RELATED (ATR), respectively (Maréchal and Zou, 2013). In animal and yeast cells, when the progression of DNA replicative polymerases is hindered, REPLICATION PROTEIN A (RPA)-coated single-stranded DNA accumulates, resulting in the recruitment and activation of ATR (Saldivar et al., 2017) through mechanisms that appear to be conserved in plants (Sweeney et al., 2009). By contrast, downstream signaling events differ between plants and animals. In plants, ATR is thought to directly phosphorylate the central DDR transcriptional regulator SUPPRESSOR OF GAMMA-RESPONSE 1 (SOG1), which in turns activates DNA repair genes and negative regulators of cell cycle progression, such as WEE1 (Preuss and Britt, 2003; De Schutter et al., 2007; Sjogren et al., 2015; Bourbousse et al., 2018). However, when treated with hydroxyurea (HU), which triggers replication stress by depleting the intracellular dNTP pool, *wee1 sog1* double mutants display a stronger growth inhibition than the corresponding single mutants, indicating that SOG1 and WEE1 function partially independently to control the replication stress response (Hu et al., 2015). Likewise, hypomorphic mutants of the replicative DNA polymerase ε (Polε) catalytic subunit POL2A show a constitutive activation of the replication stress response that is only partially dependent on SOG1. Indeed, ATR and WEE1 are crucial for the survival of *pol2a*, but the *pol2a sog1* double mutant is viable, and still shows activation of a subset of DDR genes (Pedroza-Garcia et al., 2017), indicating that part of the transcriptional response evoked by replication stress is controlled by a yet unidentified transcription factor.

Seen their role as transcriptional activators of genes required for S-phase progression, possible contributors to the transcriptional reprograming induced by replication stress could be E2F transcription factors. Among other functions, the E2Fs–RBR1 (RETINOBLASTOMA RELATED 1) module plays a well-known role in the control of DNA replication (Müller et al., 2001; Vlieghe et al., 2003; Vandepoele et al., 2005; Naouar et al., 2009). When a plant cell receives mitogenic cues, D-type cyclin-activated cyclin-dependent kinases phosphorylate RBR1, which unleashes E2F activity. In *Arabidopsis*, the E2F family comprises six members which are E2FA, E2FB, E2FC, DEL1/E2Fe, DEL2/E2Fd, and DEL3/E2Ff (Vandepoele et al., 2002). They are categorized into canonical E2Fs (E2FA, E2FB and E2FC) that function as heterodimers with their dimerization partners DPA and DPB, and non-canonical E2Fs (DEL1/E2Fe, DEL2/E2Fd and DEL3/E2Ff) that operate independently of dimerization partners (Mariconti et al., 2002; Lammens et al., 2009). Canonical E2FA and E2FB are considered as transcriptional activators because they contain a transactivation domain and stimulate S-phase entry, whereas E2FC is generally considered as a repressor (Mariconti et al., 2002; del Pozo et al., 2002; De Veylder et al., 2002; Sozzani et al., 2006; Lammens et al., 2009). E2FA and E2FB are thought to be partially redundant, because single mutants of *E2FA* and *E2FB* show no dramatic phenotypes (Yao et al., 2018; Őszi et al., 2020), whereas a double mutant is lethal (Li et al., 2017). However, some differences exist between E2FA and E2FB. For example, E2FA and E2FB play different roles in the growth inhibition triggered by UV-B exposure (Gómez et al., 2022). In addition, E2FB, together with E2FC and RBR1, was found to be a part of DREAM (DP, Rb-like, E2F, and MuvB) complexes, which are crucial for the timely succession of transcriptional waves involved in cell cycle progression and/or onset of cell differentiation (Magyar et al., 2016; Lang et al., 2021). By contrast, E2FA is not copurified with DREAM complex subunits from plant cell extracts, although it can interact with some components in the yeast two-hybrid system, suggesting that it differs from E2FB in the strength of its association to DREAM complexes (Lang et al., 2021).

In addition to their role as cell cycle regulators, several lines of evidence indicate that E2Fs control the cellular response to DNA damage and replication stress. In mammals, *E2F1* transcription is usually inactivated at the start of the S-phase by the induction of the repressive E2F6 protein, which is an E2F1 target (Giangrande et al., 2004). However, under DNA damaging conditions, the E2F1 protein is stabilized by ATM-or ATR-dependent phosphorylation (Lin et al., 2001) and accumulates at the sites of DSBs (Biswas and Johnson, 2012). Additionally, during replication stress, the E2F6 repressor is inactivated, causing sustained *E2F1* transcription that is necessary for the arrest and stabilization of replication forks and in this way prevents DNA damage (Bertoli et al., 2013; Bertoli et al., 2016). Remarkably, in cells with an impaired checkpoint control, the sustained transcription of *E2F1* is sufficient to alleviate DNA damage levels (Bertoli et al., 2016). Likewise, in tobacco BY-2 cells, the *NtE2F* gene is induced in response to high doses of UV-C (Lincker et al., 2004) and its protein localizes in distinct chromatin foci upon DNA damage (Lang et al., 2012), hinting at a possible conservation of the role of E2F transcription factors in the DDR in plants. More recently, it was found that upon DNA damage, RBR1 also colocalizes with γH2AX, a histone variant that is phosphorylated by ATM and ATR and forms foci delineating breaks in the DNA (Friesner et al., 2005), and that this is necessary for correct localization of RAD51 foci (Biedermann et al., 2017). Using chemical inhibitors of ATM and ATR, it was also found that the formation of foci of E2FA and RBR1 is dependent on both kinases (Horvath et al., 2017), recruiting the DNA repair protein BRCA1 to these foci. Furthermore, E2FB was shown to be required for cell cycle arrest induced by the crosslinking agent cisplatin (Lang et al., 2021). Collectively, these results provide strong evidence for the role of E2FA and E2FB in the cellular response to DSBs, although their contribution to the replication stress response has never been explored.

In this study, we studied the involvement of E2FA and E2FB in response to replication stress. Surprisingly, we show that E2FA and E2FB share numerous targets with SOG1. Further, we show that E2FB, rather than E2FA, activity is required to allow cell cycle progression despite replication stress. In-depth analysis of the expression behavior of common SOG1 and E2F targets demonstrates that E2FA and E2FB play both complementary and distinct roles in this pathway.

## Results

### E2FA and E2FB share targets with SOG1

E2FA, E2FB and RBR1 have all recently been shown to play a role in the plant’s DSB response (Biedermann et al., 2017; Horvath et al., 2017; Lang et al., 2021). Exploiting our recent analysis of E2FA and E2FB targets (Gombos et al., 2022), we investigated whether E2FA/B shared common targets with SOG1. When comparing the target genes of E2FA (Supplemental Table S1A), E2FB (Supplemental Table S1B) and SOG1 as determined by tandem chromatin affinity purification (TChAP) (Verkest et al., 2014) or chromatin immunoprecipitation followed by sequencing (ChIP-seq) experiments (Bourbousse et al., 2018; Gombos et al., 2022), we found that a greater number of SOG1 target genes was also targeted by at least one of the two E2F transcription factors than what would be expected by chance (Figure 1A, Supplemental Table S1C). Among these, the *WEE1* gene could be found, encoding a cell cycle inhibitory kinase implicated in the replication stress response (De Schutter et al., 2007). Interestingly, residual activation of the *WEE1* promoter was observed in response to HU-induced replication stress in a *sog1* mutant and required the E2F-binding site found in its promoter, confirming a potential involvement of E2FA/B in the replication stress response (Supplemental Figure S1). We next compared the position of experimentally identified E2FA- and E2FB-binding sites with those previously identified for SOG1 (Bourbousse et al., 2018). Both E2FA and E2FB bound the common target genes at positions close to the SOG1-binding site (Figure 1B-E, Supplemental Table S1C), suggesting that E2FA/B and SOG1 bind in close proximity to each other on their target promoters. The observation that E2FA/B can activate *WEE1* in response to HU independently of SOG1 (Supplemental Figure S1) and the significant overlap between putative E2FA/B and SOG1 target genes (Figure 1A) prompted us to test whether E2FA/B play a role in the replication stress response.

**Figure 1:**
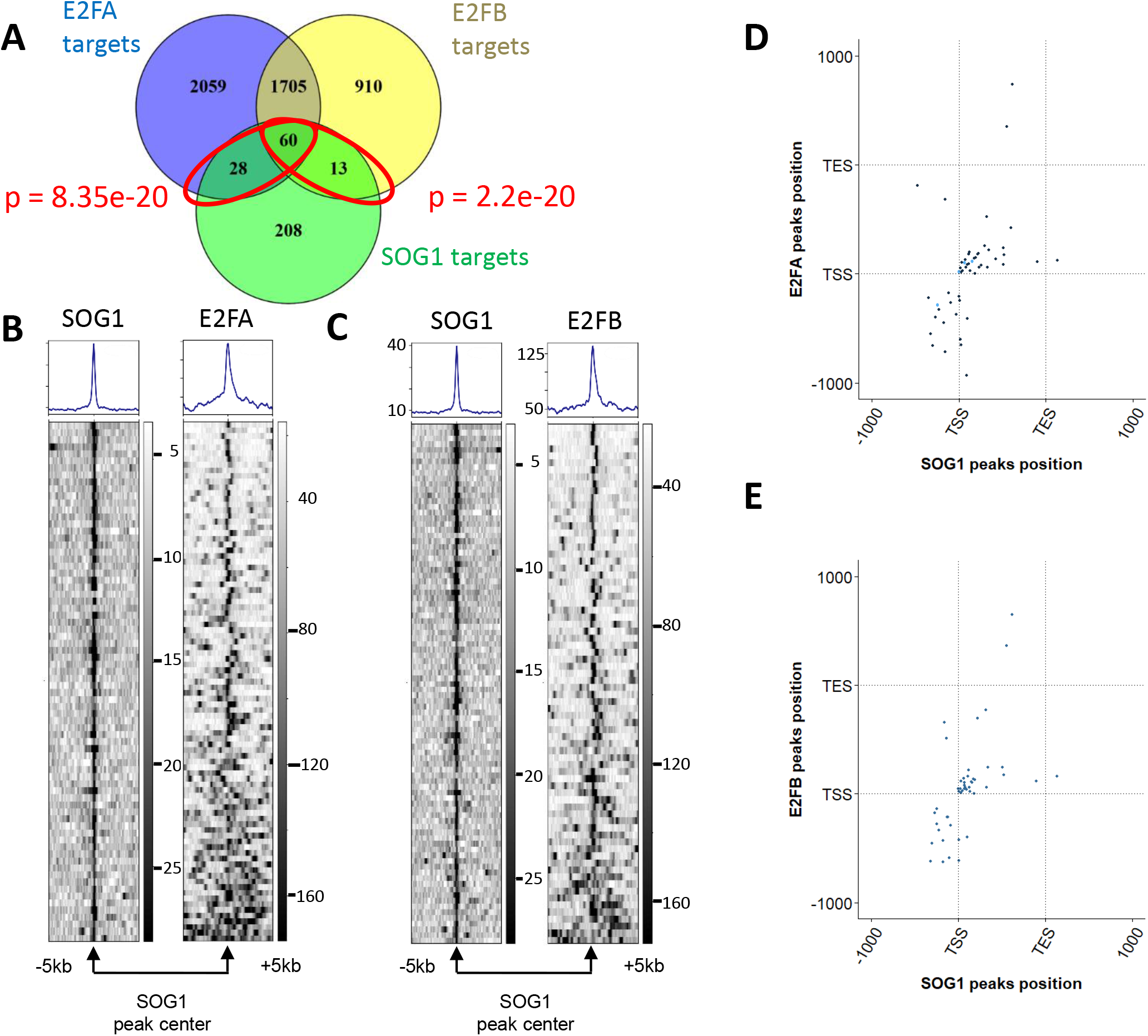
E2FA, E2FB and SOG1 share common target genes. A: Venn diagram showing the overlap between SOG1 (Bourbousse et al., 2018), E2FA (as defined by the union of targets found by TChAP (Verkest et al., 2014) and by ChIP-seq (Gombos et al., 2022; FDR < 0.05 and enrichment > 2.4) and E2Fb target genes (Gombos et al., 2022; FDR < 0.05, enrichment > 2). Significance of overlap was estimated with Fisher’s exact test (total number of loci: 38,194 according to Araport11, Cheng et al., 2017). B: Heatmaps showing E2FA and SOG1 binding on their common target genes, centered around the SOG1 binding site. SOG1 and E2FA binding sites are at very similar positions on most of their common targets, as illustrated by the metaplot above the heatmap. C: Heatmaps showing E2FB and SOG1 binding on their common target genes, centered around the SOG1 binding site. SOG1 and E2FB binding sites are at very similar positions on most of their common targets, as illustrated by the metaplot above the heatmap. D, E: Density plots showing overlap of SOG1 and E2FA (D) or E2FB (E) using hexagonal binning routine on their common target genes. Each dot represents the distance from the peak midpoint to the nearest gene. The *y*-axis shows the location of the E2FA (D) or E2FB (E) peak midpoint compared with gene position, while the *x*-axis indicates the position of the SOG1 peak midpoint relative to the nearest gene. Most dots occur close to the diagonal of the graph, showing that E2FA/B and SOG1 bind neighboring sequences. TSS: transcription start site, TES: transcription end site.

### Loss of E2FB strongly aggravates growth defects triggered by replication stress

To explore the role of E2FA and E2FB in the replication stress response, we used the hypomorphic mutant for DNA polymerase ε (Polε), *pol2a-4* (hereafter referred to as *pol2a*) that shows constitutive replication stress (Pedroza-Garcia et al., 2017), and generated double and triple mutant combinations between *pol2a, sog1* and *e2fa* or *e2fb* mutants. Two independent T-DNA insertion lines have been described for both *E2FA* and *E2FB* (Berckmans et al., 2011a; Berckmans et al., 2011b). In the case of *E2FA*, the *e2fa-1* allele appears to be a null mutant lacking E2FA protein accumulation, whereas *e2fa-2* accumulates significant levels of a truncated protein (Leviczky et al., 2019). In the case of *E2FB*, the protein cannot be detected in protein extracts of either *e2fb-1* or *e2fb-2* mutants (Leviczky et al., 2019). In terms of protein function, viable *e2fa e2fb* double mutants have been obtained using the *e2fa-2* but not the *e2fa-1* allele (Heyman et al., 2011), suggesting that the truncated protein accumulating in *e2fa-2* mutants is at least partially functional. For our genetic analysis, we therefore used only the *e2fa-1* mutant allele, in which *E2FA* loss of function is likely full, and both *e2fb* alleles. Throughout the manuscript, we show results obtained for the *e2fb-1* allele, but the *e2fb-2* allele systematically gave the same results. Phenotypically, *e2fa-1* and *e2fb-1* single mutants did not show a growth reduction compared with the wild type (Col0), whereas *pol2a* mutants were significantly smaller (Figure 2A, B). Growth reduction was more severe in the *pol2a sog1* double mutant, consistent with the hypersensitivity of the *sog1* mutant to replication stress (Pedroza-Garcia et al., 2017). The *pol2a e2fa-1* mutant rosette size was identical to that of the *pol2a* parent, whereas the *e2fb-1 pol2a* mutant was slightly smaller. Strikingly, the *e2fb-1 pol2a sog1* triple mutant showed a more severe growth defect than *pol2a sog1*, a phenomenon not observed with the *e2fa-1* mutation (Figure 2A, B). We also analyzed the root length of the various mutants, and observed that E2FB, but not E2FA, is required for root growth in plants suffering from constitutive replication stress (Figure 2C), particularly in the absence of SOG1. These data show that E2FB contributes to the plant’s response to replication stress, allowing growth maintenance in spite of the replication defects.

**Figure 2:**
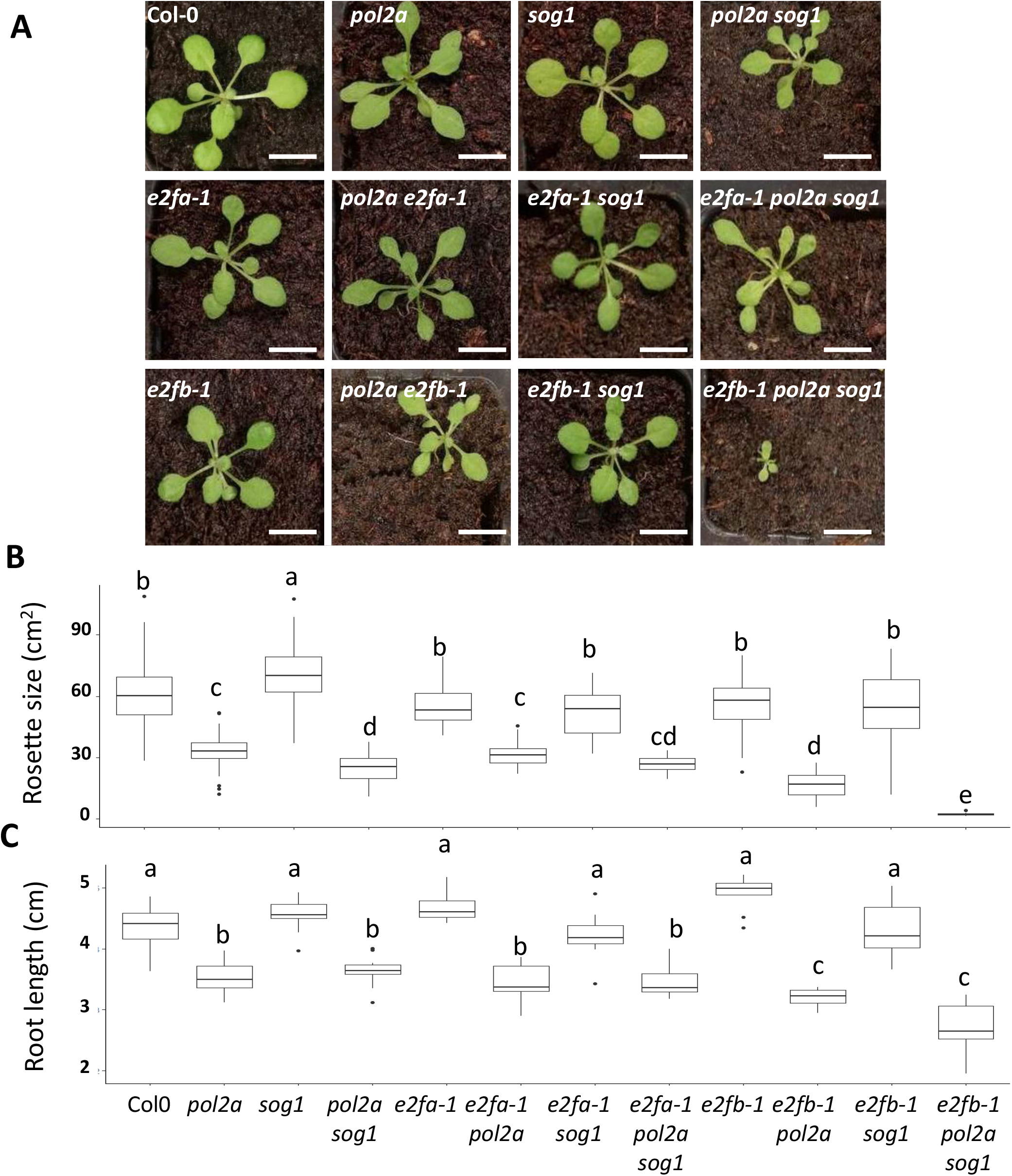
E2FB, but not E2FA, is required for sustained plant growth in response to replication stress. A: Representative rosette phenotype of 30-day-old plants of the indicated genotype. Bars = 1cm. B-C: Quantification of rosette area (B) or root length (C) in the indicated genotypes. Data are mean ± SD from at least 15 (B) or 20 (C) measurements for each line and are representative of two independent experiments. On box plots, the vertical size of the boxes shows the interquartile range, and the whiskers correspond to 1.5x the IQR. The horizontal line corresponds to the median. Individual dots indicate values falling outside of this range. Significant differences from the wild type are determined by one-way ANOVA with post-hoc Tukey HSD; *, p < 0.05. Different letters indicate statistically significant differences (ANOVA and Tukey test p < 0.05 for B or p < 0.01 for C).

### E2FB positively regulates meristem size and cell cycle progression in response to replication stress

The severe growth reduction observed in *e2fb-1 pol2a sog1* triple mutants, and to a lesser extent, in *e2fb-1 pol2a* double mutants, likely results from cell proliferation defects. To test this hypothesis, we first measured the root meristem size in all genotypes. As shown in Figure 3, replication stress triggered by Polε deficiency resulted in a reduced meristem size. Whereas this defect was not significantly aggravated in the absence of SOG1, root meristem length was further reduced in *ef2b-1 pol2a* and *e2fb-1 pol2a sog1* mutants (Figure 3B). Again, this effect was not observed in *e2fa-1 pol2a* and *e2fa-1 pol2a sog1* mutants (Supplemental Figure S2). These results confirmed that E2FB, but not E2FA, plays a crucial role in proliferating cells to protect them from cell proliferation arrest triggered by replication stress.

**Figure 3:**
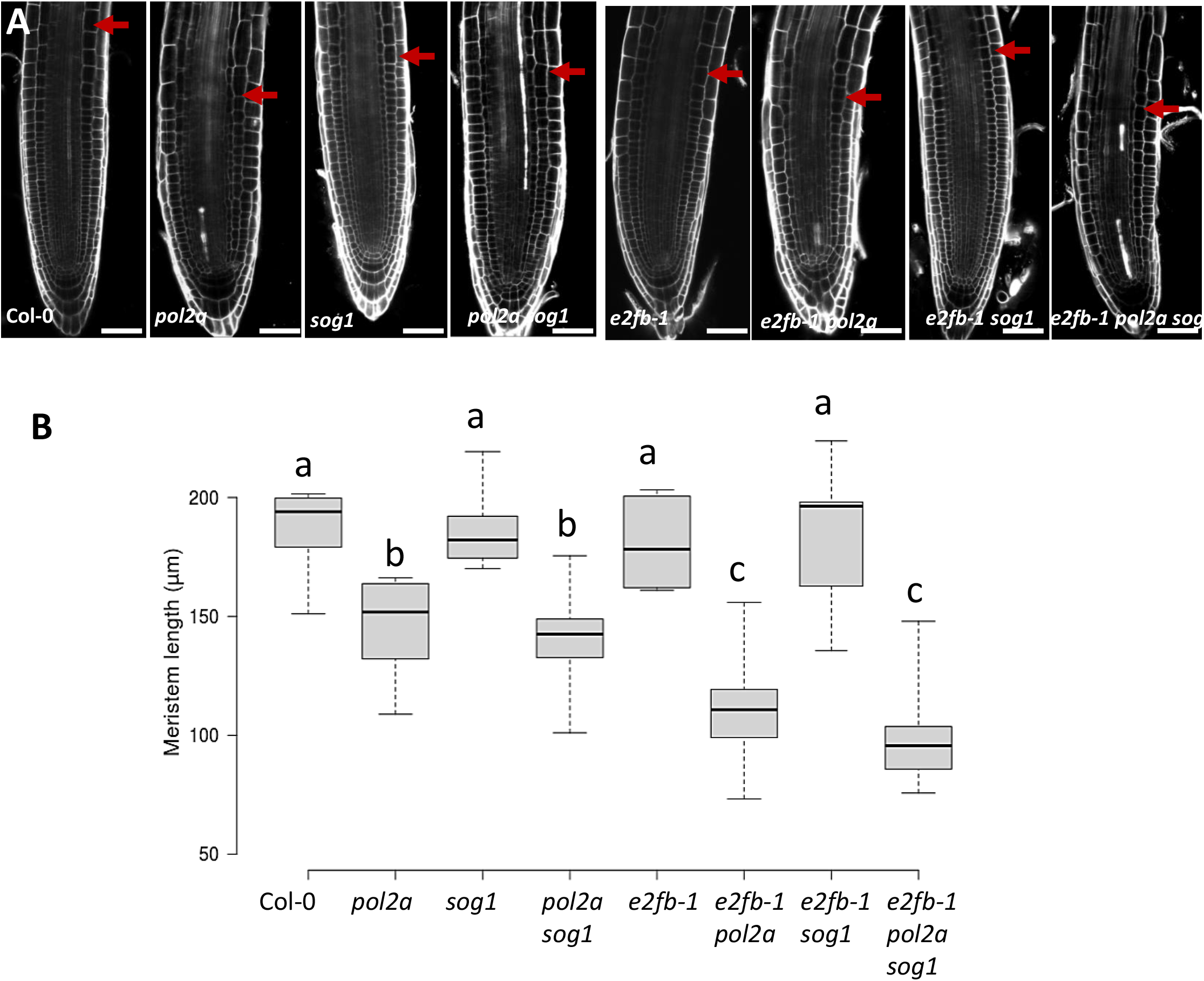
Loss of E2FB further reduces root apical meristem length in the *pol2a* background. A: Representative confocal images of 7-day-old root apical meristem of wild type (Col-0), *pol2a, sog1, pol2a sog1, e2fb-1, e2fb-1 pol2a, e2fb-1 sog1* and *e2fb-1 pol2a sog1*. Cell walls were stained with propidium iodide (PI). Red arrows indicate the limit of the apical root meristem. Bars = 20 μm. B: Quantification of root meristem length in indicated genotypes (n > 10). On box plots, the vertical size of the boxes shows the interquartile range, and the whiskers correspond to 1.5x the IQR. The horizontal line corresponds to the median. Individual dots indicate values falling outside of this range. Different letters indicate statistically relevant differences (ANOVA followed by Tukey test p < 0.05). Data are representative of two independent experiments.

To further dissect how E2FB affects cell proliferation in response to replication stress, we analyzed cell cycle progression into more detail in all mutant combinations. We first analyzed the distribution of cell cycle phases of nuclei in flower buds. The proportion of nuclei in each cell cycle phase was the same in the wild type and *sog1*, single *e2fb-1* and double *e2fb-1 sog1* mutants (Figure 4A). The proportion of S-phase cells increased in all mutant combinations containing the *pol2a* mutation, consistent with our previous findings (Pedroza-Garcia et al., 2017). In addition, we observed that the proportion of G2 nuclei was increased at the expense of the proportion of G1 nuclei in *e2fb-1 pol2a* and *e2fb-1 pol2a sog1* mutants compared to *pol2a* and *pol2a sog1* mutants, respectively (Figure 4A). This phenomenon was not observed in *e2fa-1* mutant combinations (Supplemental Figure S3). These data suggest that E2FB positively regulates cell cycle progression through G2 and the onset of the G2/M transition in replicative-stress exposed cells.

**Figure 4:**
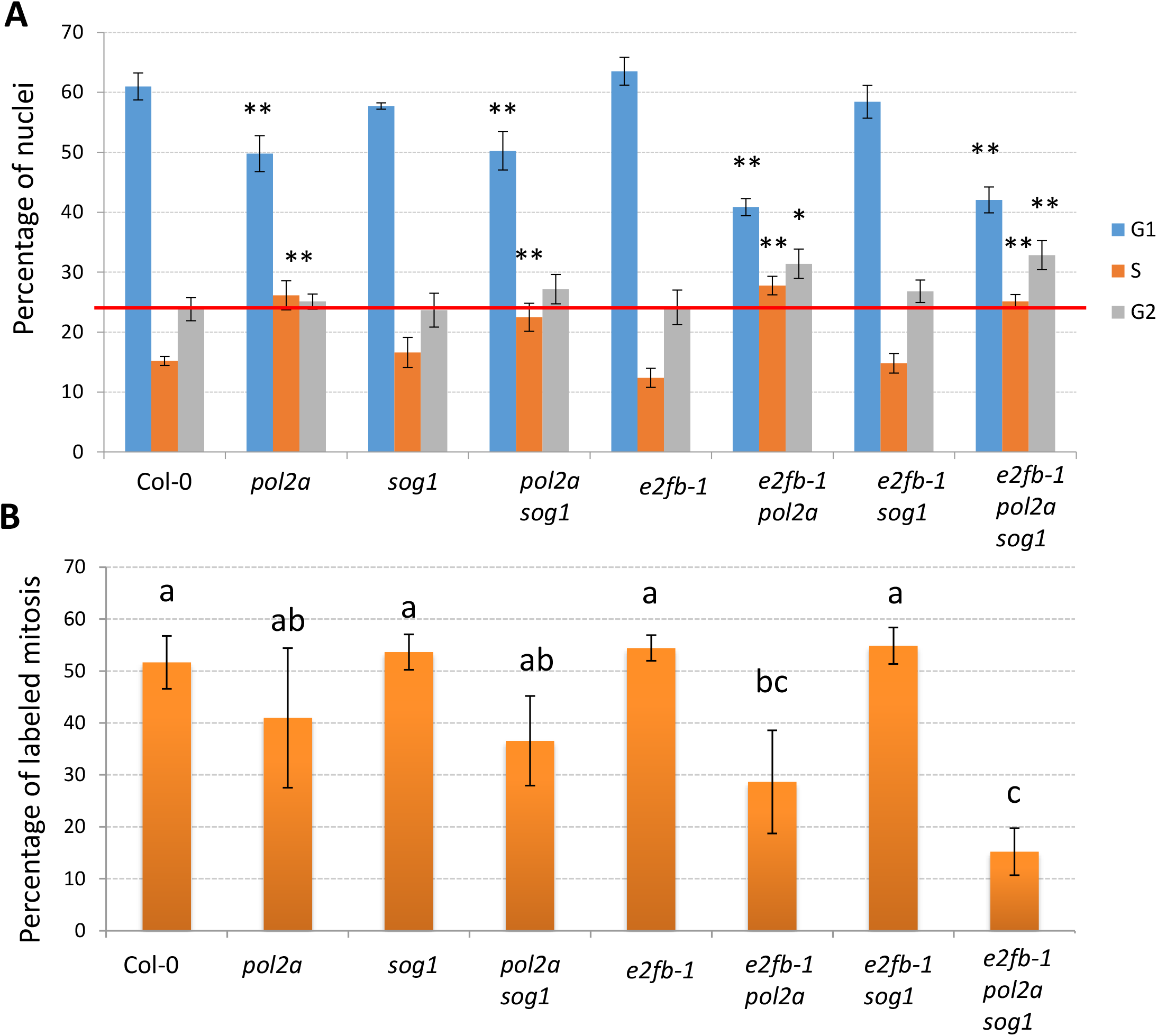
E2FB positively regulates G2/M progression in response to replication stress. A: Flow cytometry was used to analyze the cell cycle phases in the flower buds of Col-0, *pol2a, sog1, pol2a sog1, e2fb-1, e2fb-1 pol2a, e2fb-1 sog1* and *e2fb-1 pol2a sog1* mutant lines. For each cell cycle phase, the percentage of nuclei was calculated. Values are average ± SD. Asterisks denote significant differences with respect to percentages observed in the wild type (Kruskal-Wallis test, * p < 0.05, ** p < 0.01). Data are representative of three biological replicates. The red line indicates the proportion of G2 nuclei in the wild-type. B: Mitosis in five-day-old roots of all mutant lines labeled with ethynyl-2’-deoxyuridine (EdU). The bar graph represents the percentage of labeled mitosis in the indicated genotypes 5 h after transfer on EdU-containing medium. Data represent at least 10 roots and are representative of three independent experiments. Different letters indicate statistically significant differences (Kruskal-Wallis test).

To confirm the flow cytometry result, we next performed cumulative ethynyl-2’-deoxyuridine (EdU) labeling on four-day-old seedlings. EdU was incorporated extremely slowly in *pol2a sog1 e2fb-1* triple mutants (Supplemental Figure S4), which made it difficult to robustly estimate cell cycle or S-phase length as described before (Yin et al., 2014). As an alternative, we counted the proportion of roots showing EdU-labeled mitosis 5 h after EdU application, to monitor how S and G2 progression was affected in the various mutant combinations. As expected, the proportion of labeled mitosis was reduced in *pol2a* mutants compared with the wild type, whereas it was unchanged in *sog1, e2fb-1* or *e2fb-1 sog1* mutants (Figure 4B). Furthermore, *e2fb-1 pol2a* double and *e2fb-1 pol2a sog1* triple mutants showed a lower percentage of labeled mitosis compared with *pol2a* and *pol2a sog1* mutants, respectively (Figure 4B). Together, these results confirmed that upon replication stress, E2FB is required for cell cycle progression through G2, and that this function is particularly important in the absence of SOG1, suggesting that SOG1 and E2FB may act in parallel to maintain the proliferative capacity in replication stress-exposed cells.

### E2FB and SOG1 both regulate the expression of replicative stress-induced genes

To gain insight into the mechanisms underlying the role of E2FB in the replication stress response, we compared gene expression changes in shoot apices of *pol2a, pol2a sog1* and *e2fb-1 pol2a sog1* mutants (Supplemental Table S2). Compared with wild type plants, we found 1,822, 2,599 and 3,512 upregulated genes in *pol2a, pol2a sog1* and *e2fb-1 pol2a sog1* mutants, respectively (FDR < 0.05). Among those, 1,095 were commonly upregulated in all three mutant lines (Figure 5A, Supplemental Table S3). Gene Ontology (GO) analysis of these 1,095 genes revealed a significant enrichment in GO terms such as DNA repair, DNA replication, and negative regulation of the cell cycle (Figure 5B), consistent with the constitutive replication stress triggered by *POL2A* deficiency and induced cell cycle defects (Pedroza-Garcia et al., 2017). As anticipated, genes specifically upregulated in the *e2fb-1 pol2a sog1* mutant were not enriched in E2F target genes, indicating that most of them are likely indirectly regulated by E2Fs.

**Figure 5:**
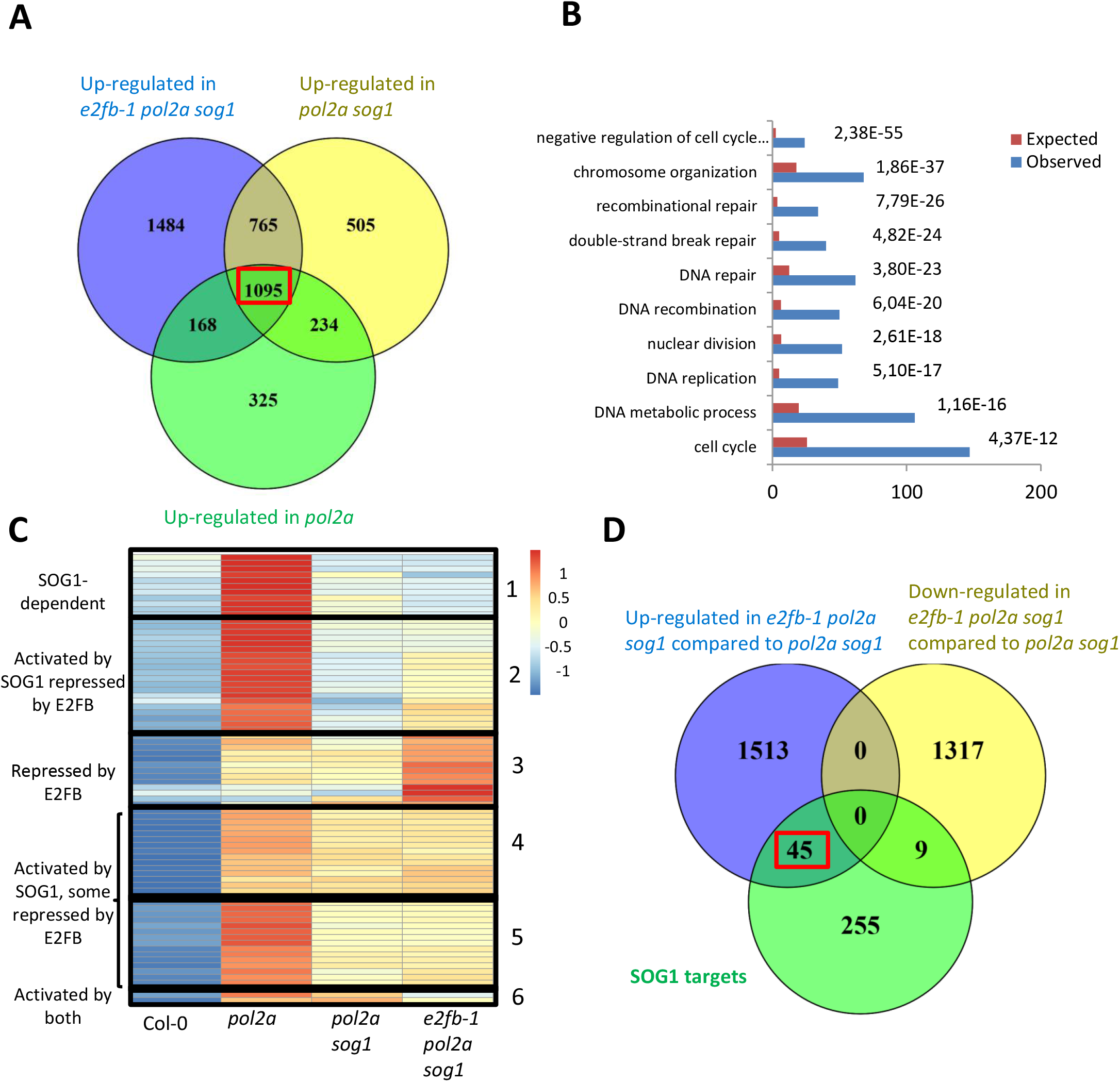
E2FB and SOG1 cooperate to control replicative stress-induced transcriptional changes. A: Venn diagram showing the overlap between upregulated genes in *pol2a, pol2a sog1* and *e2fb-1 pol2a sog1* compared with wild-type plantlets in the shoot apex. Apices were collected from 7-day-old plantlets by removing cotyledons and hypocotyls. B: GO-term analysis of genes upregulated in all mutant lines. C: K-means clustering of expression changes of SOG1 target genes in the different mutant lines. The analysis was performed using genes that were upregulated in at least one mutant. D: Overlap between SOG1 target genes and mis-regulated genes in triple mutants compared with *pol2a sog1* double mutants. SOG1 targets are significantly more represented amongst upregulated genes, indicating that E2FB acts as a repressor of these genes.

To understand the behavior of SOG1 target genes in the different genotypes, we performed a k-means clustering analysis of the expression levels of SOG1 targets in *pol2a, pol2a sog1* and *e2fb-1 pol2a sog1* mutants (Figure 5C, Supplemental Table S4A). Out of the 309 SOG1 targets, 78 were significantly upregulated (FDR < 0.05, log_2_FC > 0.5) in at least one genotype compared with the wild type and were kept in the analysis. This approach unraveled six gene clusters that displayed contrasting expression profiles. Genes in cluster 1 appeared to be controlled by SOG1 only, as they are upregulated in *pol2a* mutants but not in *pol2a sog1* or *e2fb-1 pol2a sog1* mutants and are not enriched in E2F targets (Supplemental Table S4B). Conversely, genes in cluster 3 were affected mainly by the *e2fb-1* mutation, as they are only slightly upregulated in *pol2a* and *pol2a sog1* mutants and very strongly upregulated in the triple mutant. Genes in the remaining clusters were affected by both E2FB and SOG1. Clusters 4 and 5 comprised genes that were upregulated in all three mutant lines, although this induction was less pronounced in the *sog1* mutant background. Among those, some were expressed at similar levels in *pol2a sog1* and *e2fb-1 pol2a sog1* mutants, whereas others were upregulated in the absence of E2FB. Likewise, genes in cluster 2 were downregulated in *pol2a sog1* mutants compared with the *pol2a* single mutant, but their expression level increased in the triple mutant, suggesting that these SOG1 target genes were under the antagonistic control of E2FB and SOG1. Interestingly, clusters 2, 4 and 5 were particularly enriched in E2F targets (Supplemental Table S4B), further supporting the hypothesis that they are under dual control of SOG1 and E2FB. Indeed, SOG1 targets were significantly enriched amongst genes upregulated in the *e2fb-1 pol2a sog1* triple mutant compared with the *pol2a sog1* mutant (p =1.118e-5, Fisher’s exact test, total number of genes detected in the RNA-Seq analysis = 20,946, Figure 5D), further confirming that *bona fide* SOG1 targets can be activated independently of SOG1 in response to replication stress.

Among shared E2FB/SOG targets that appeared to be antagonistically controlled by SOG1 and E2FB, we found the *ANAC044* and *ANAC085* transcription factor genes (respectively found in clusters 4 and 2) that have been shown to negatively regulate G2/M progression. This result was confirmed by RT-qPCR (Supplemental Figure S5A). Interestingly, *ANAC044* and *ANAC085* are not only direct SOG1 and E2FB targets, but also RBR1 targets ((Gombos et al., 2022), Supplemental Figure S5B, C), suggesting that E2FB could repress them through its interaction with RBR1.

Together, our transcriptomics results suggest that SOG1 and E2FB have mostly antagonistic effects on their common targets, and that E2FB could be important to mitigate the induction of cell cycle arrest genes induced by SOG1. This hypothesis raises the question of the molecular mechanism allowing the activation of SOG1 targets in the absence of both SOG1 and E2FB. Interestingly, GO analysis of all genes that were specifically upregulated in *pol2a sog1* and *e2fb-1pol2a sog1* mutants and not in *pol2a* single mutants compared with wild-type plants revealed a significant enrichment in terms of related to cell cycle regulation (Supplemental Figure S6A). In addition, E2FA and E2FB targets were significantly enriched in this list (p = 3.365e-04 and p = 1.734e-05 respectively, Fisher’s exact test, total number of genes detected in the RNA-Seq analysis = 20,946, Supplemental Figure S6B). It is worth noting that E2F targets were also highly enriched among genes that were commonly upregulated in all three mutant lines (428 out of 1,095, p = 8.179e-36, Fisher’s exact test, total number of genes detected in the RNA-Seq analysis = 20,946). Together, these results indicate that other E2Fs, and notably E2FA, could be involved in the regulation of these genes.

### E2FA, E2FB and SOG1 cooperate to allow cell cycle progression in response to replication stress

To test the possibility that E2FA could contribute to the replication stress response in the absence of E2FB and SOG1, we used the partial loss-of-function allele of E2FA (*e2fa-2*), and generated *e2fa-2 e2fb-1 sog1* triple mutants. To induce replication stress, plantlets were exposed to the replication inhibitor HU. As published before, *sog1* mutants are slightly sensitive to replication stress (Hu et al., 2015), whereas *e2fa, e2fb* and *e2fa-2 e2fb-1* showed no such hypersensitivity (Figure 6). Both *e2fa* and *e2fb* mutant alleles behaved in the same way. Similarly, as observed in response to constitutive replication stress induced by the *pol2a* mutant (Figure 2B), *e2fb-1 sog1* but not *e2fa-1 sog1* double mutants were more sensitive to HU than *sog1* mutants, consistent with the hypothesis that E2FB plays a more prominent role than E2FA in the replication stress response. Strikingly, the *e2fa-2 e2fb-1 sog1* triple mutant displayed even stronger HU hypersensitivity, with root growth being almost completely blocked after transfer on HU (Figure 6), and cell death being induced in the root meristem (Supplemental Figure S7), which reveals a contribution of E2FA to the replication stress response when both E2FB and SOG1 are inactivated.

**Figure 6:**
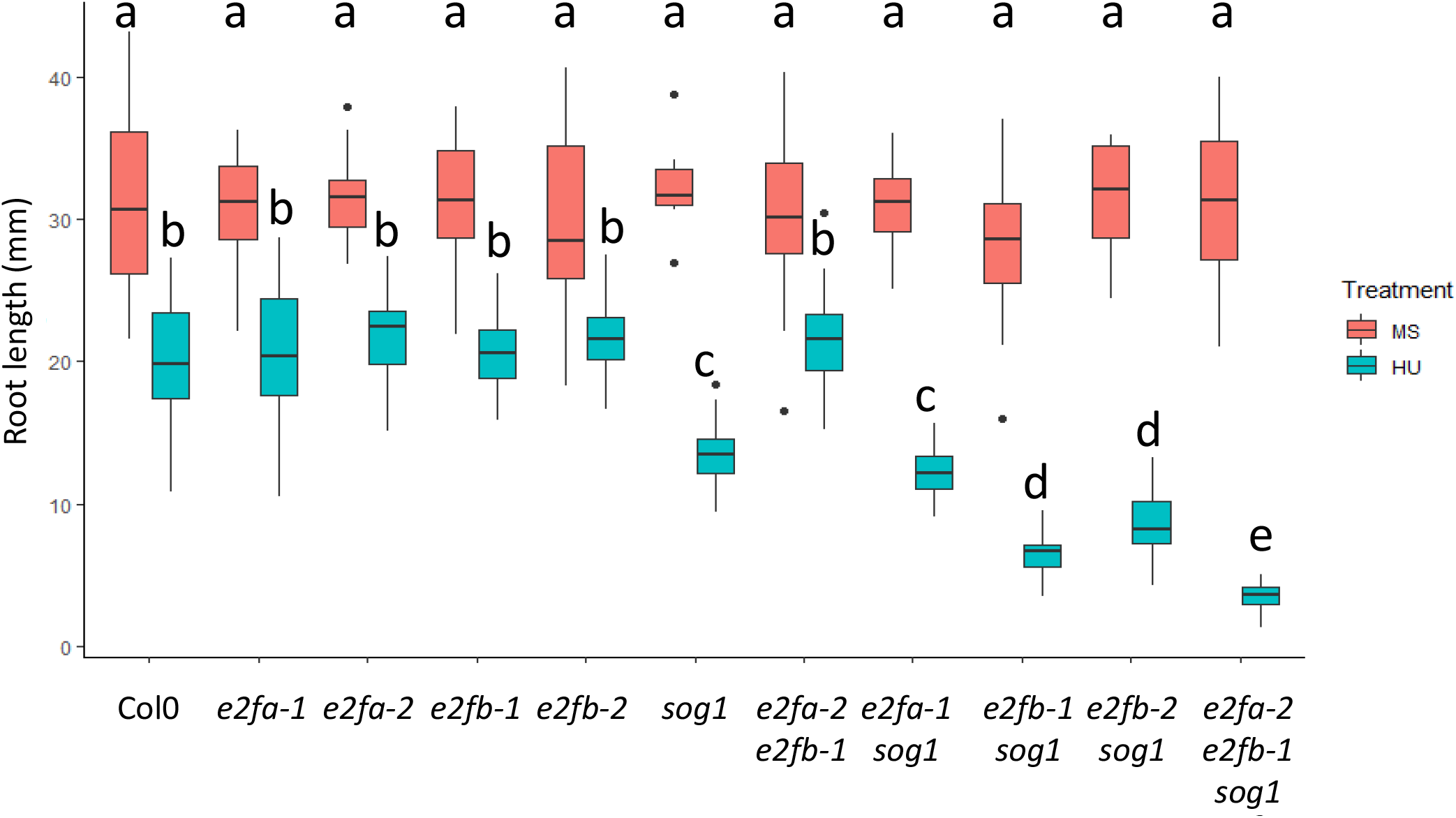
Simultaneous loss of E2FA, E2FB and SOG1 abolishes the plant’s ability to tolerate replication stress. Plantlets were germinated on 0.5x MS, and transferred on medium supplemented with 1 mM HU after five days. Root lengths were measured after ten days. Data presented are mean ± SD (n > 20). Significant differences from the wild type are determined by one-way ANOVA with post-hoc Tukey HSD; *, p < 0.05. Different letters indicate statistically significant differences (ANOVA and Tukey test p < 0.01).

Finally, because hindrance of fork progression is inevitable during the S-phase, we investigated whether E2F deficiency may trigger replication stress even in a wild-type background. Indeed, *e2fa-2 e2fb-1* double mutants (but not the single mutants) have been shown to display reduced root meristem size and a global slight reduction in cell proliferation (Gómez et al., 2022) (Supplemental Figure S8). Although this phenotype could be due to the role of E2Fs as activators of cell cycle progression, it may also reflect an inefficient response to basal levels of replication stress that generally happen during the S-phase. In line with this hypothesis, we found that *e2fa-2 e2fb-1* double mutants displayed a prolonged S-phase and increased total cell cycle length (Figure 7). Importantly, these defects were abolished by the *sog1* mutation, indicating that E2F deficiency triggers replication stress leading to a SOG1-dependent cell cycle delay. To confirm this hypothesis, we performed an RNA-Seq analysis on root tips of 7-day-old seedlings grown under control conditions. A total of 148 genes were found to be downregulated in the *e2fa-2 e2fb-1* mutant (FDR < 0.01), of which 73 are likely direct E2FA or E2FB target genes based on ChIP data (Figure 8A, Supplemental Table S5A, B). This list includes described components of the DNA replication machinery (Supplemental Table S6), and GO enriched categories (obtained through http://geneontology.org/), including DNA replication initiation, DNA duplex unwinding and chromatin assembly (Supplemental Table S7). Next to the 148 downregulated genes, we found 345 genes to be upregulated (FDR < 0.01, FC > 1.5) of which 145 are E2FA-or E2FB-bound genes according to ChIP data (Figure 8A, Supplemental Table S5C), consistent with recent evidence that the canonical E2FA/B play a key role in the repression of cell cycle genes, likely through their association with RBR1 (Gombos et al., 2022). GO categories enriched in the list of E2F-bound upregulated genes mainly indicate DNA repair, recombinational repair and response to gamma irradiation and X-ray (Supplemental Table S8). Quantitative RT-PCR of a selected number of genes from both down- and upregulated genes confirmed the results from the RNA-Seq (Figure 8B, C). Strikingly, upregulated *PARP1* and *SMR7* gene expression in the *e2fa-2 e2fb-1* double mutant was repressed in the *e2fa-2 e2fb-1 sog1* triple mutant. Differently to the *MCM9* gene that showed no transcriptional repression, the former two are *bona fide* SOG1 target genes, indicative that absence of E2FA and E2FB triggers a SOG1 dependent transcriptional response (Figure 8C). Accordingly, among the 345 genes upregulated in *e2fa-2 e2fb-1* double mutants, 43 were SOG1 targets (a number significantly greater than could be expected by chance, p = 7.746e-24, Fisher’s exact test, total number of genes detected in the RNA-Seq experiment = 17,028), further confirming that loss of E2FA and E2FB triggers replication stress and SOG1-dependent activation of DDR genes.

**Figure 7:**
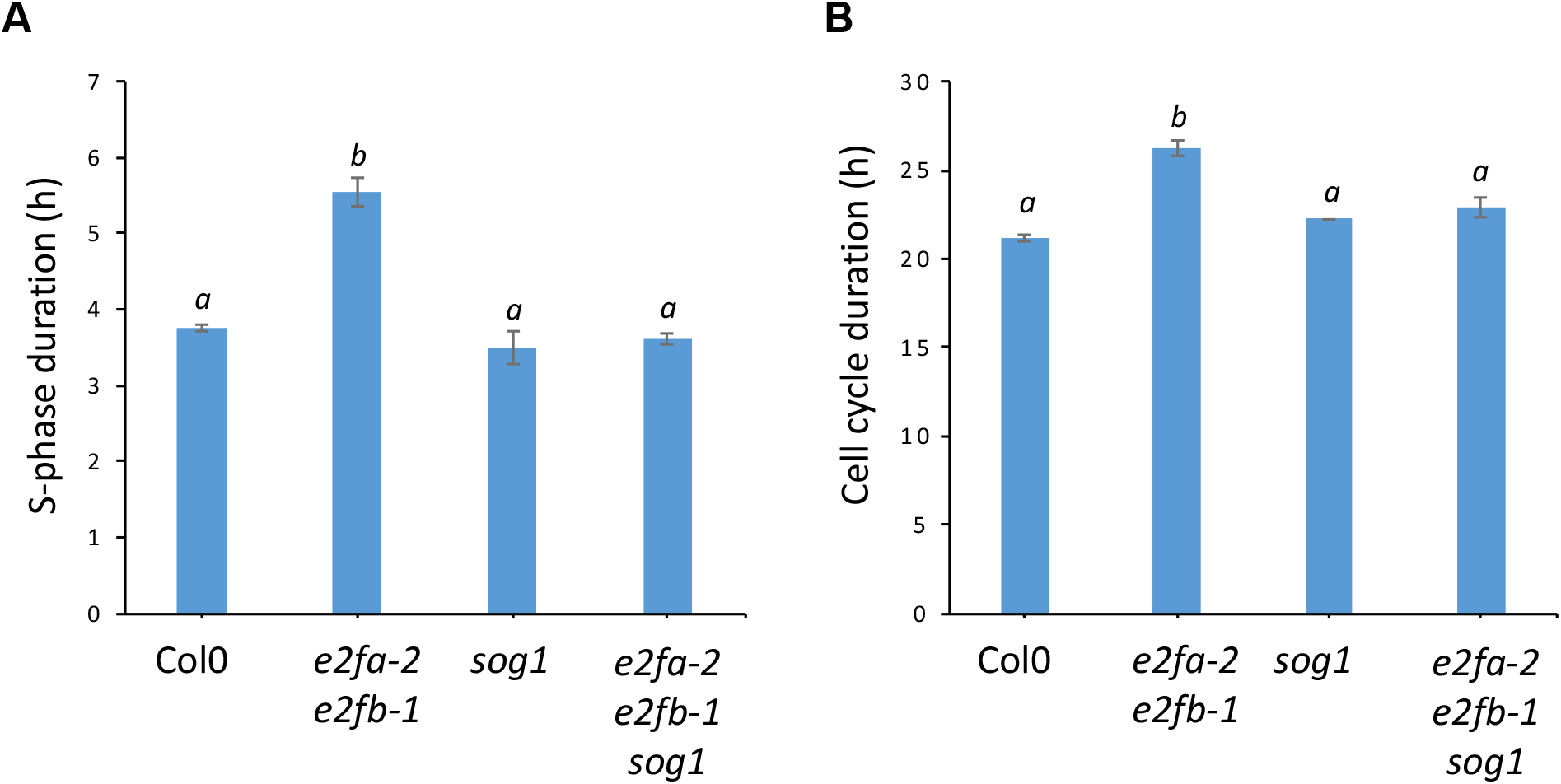
E2F deficiency triggers replication stress and SOG1-dependent cell cycle delay. A-B: S-phase (A) and total cell cycle duration (B) were measured using a time course of EdU staining according to the protocol of Hayashi et al. (2013).

**Figure 8:**
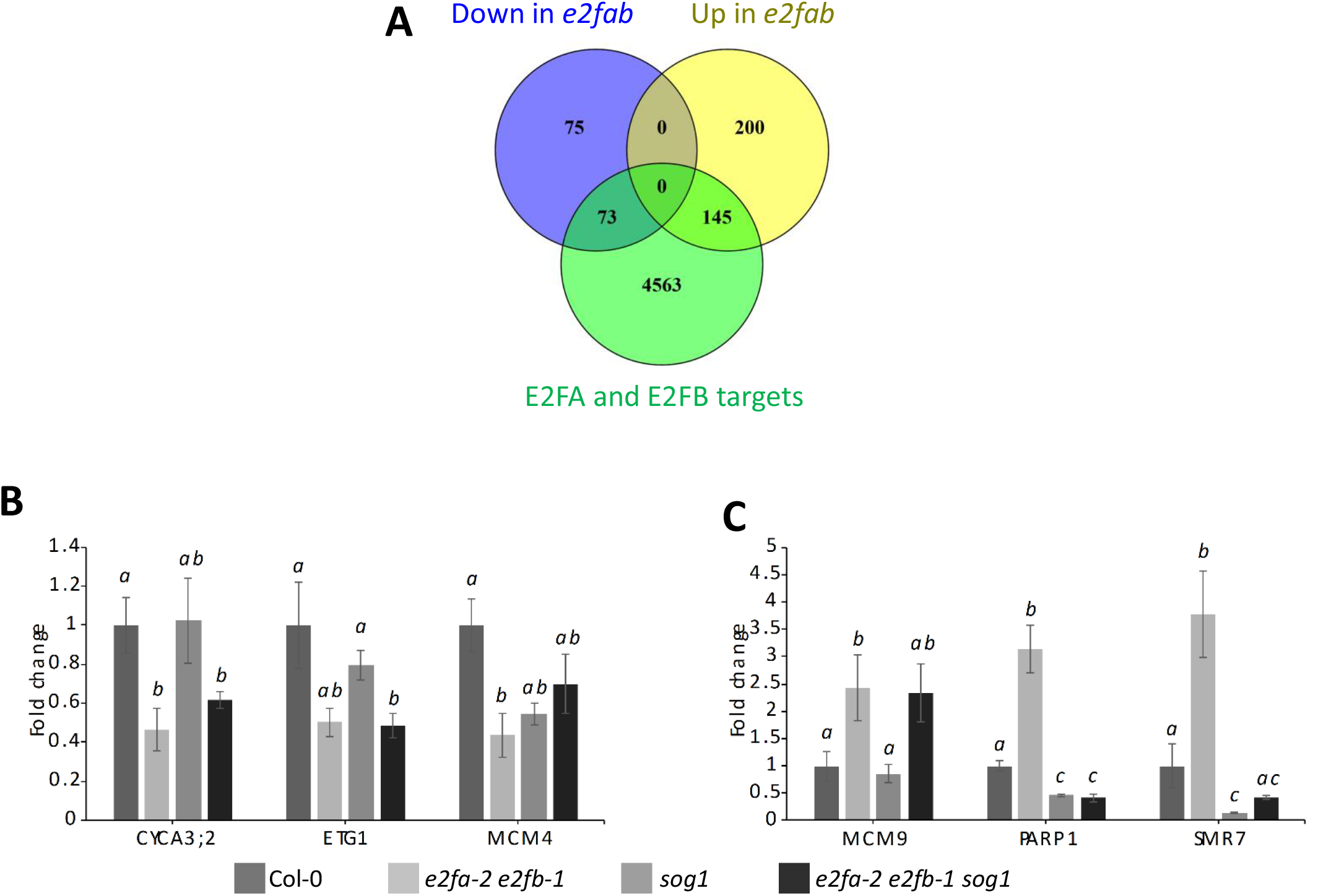
Analysis of differentially expressed genes in the *e2fa-2 e2fb-1* mutant. A: Overlap between up- and downregulated genes in *e2fa-2 e2fb-1* mutant against experimental dataset of E2FA-bound genes (Verkest et al., 2014). B-C: Relative expression levels of genes downregulated (B) or upregulated (C) in root tips of 7-day-old wild-type, *e2fa-2 e2fb-1, sog1* and *e2fa-2 e2fb-1 sog1* mutant seedlings. Data represent mean ± SEM. Experiment was done in three technical and three biological repeats of at least 100 root tips. Significance was tested with Student’s t-test. Means with different letters are significantly different (p < 0.05).

## DISCUSSION

E2Fs are core cell cycle regulators of the G1/S transition that are evolutionarily conserved over most multicellular eukaryotes, including animals and the green lineage (Bertoli et al., 2013). Although functional diversification of E2Fs has been described into details in animals (Ishida et al., 2001), our understanding of plant E2Fs’ specific functions remains limited.

### E2FB plays a more prominent role than E2FA in the plant’s response to replication stress

Based on our observation that the Arabidopsis E2FA and E2FB transcription factors share several target genes with the central DDR regulator SOG1, we investigated their contribution to the plant’s replication stress response. We found that loss of E2FB, but not of E2FA, severely aggravated the developmental defects of the *pol2a-4* mutant that suffers from constitutive replication stress (Pedroza-Garcia et al., 2017), as well as the sensitivity of the *sog1* mutant to the DNA-replication blocking drug HU. This requirement for E2FB for replication stress tolerance was particularly obvious in the *sog1* background, suggesting that E2FB and SOG1 act in parallel to cope with replication defects (Figure 9A). At the cellular level, we observed that the percentage of cells in the G2 phase was significantly higher in these lines, suggesting that the length of the G2 phase was increased by E2FB loss of function, which was confirmed by estimating the proportion of EdU-labeled mitoses over time after transfer of plantlets to EdU-containing medium. Together, our results suggest that E2FB could allow the progression of cells through G2/M under replication stress conditions.

**Figure 9:**
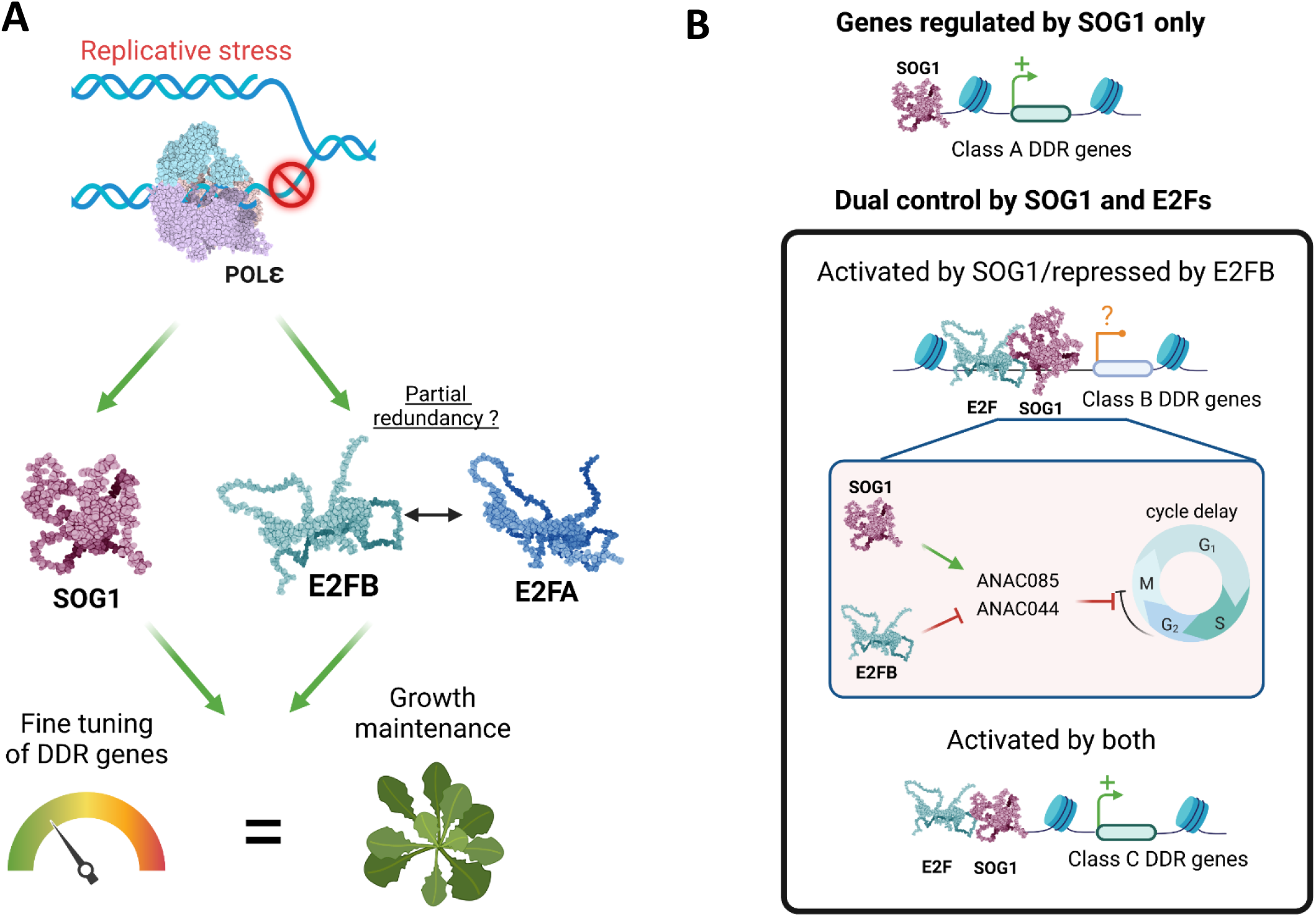
SOG1, E2FA and E2FB act on both distinct and common targets to fine-tune the plant DDR. A: Our genetic analysis shows that SOG1 and E2Fs function independantly to fine-tune DDR gene expression and allow sustained plant growth in response to replication stress. B: DDR genes can be distributed amongst three classes. One first set of genes (class A) depends only on SOG1 for their activation. Class B genes are antagonistically regulated by SOG1 and E2FB, suggesting that E2FB could dampen SOG1-dependent cell cycle arrest to avoid complete developmental arrest. Among those, negative regulators of the cell cycle such as ANAC085 and ANAC044 may contribute to the severe cell cycle arrest observed in *e2fb pol2a sog1* triple mutants. Class C genes are also targeted both by SOG1 and E2FA/B, and remain induced at similar levels by replication stress even in the absence of SOG1 and E2FB, suggesting that they are redundantly controlled by both E2FA and E2FB. Image created withy BioRender.com.

We found that negative regulators of the cell cycle such as *ANAC044* and *ANAC085* are upregulated in *e2fb-1 pol2a sog1* mutants compared with *pol2a sog1* mutants, suggesting that E2FB may allow G2 to M progression by inhibiting the repressors of the G2/M transition. Importantly, a role in allowing sustained cell proliferation in response to replication stress seems specific to E2FB, since loss of E2FA did neither alter the sensitivity of *sog1* mutants to HU, nor affect plant growth, meristem size or the proportion of G2 cells in the *pol2a* or *pol2a sog1* mutants. These nonoverlapping roles could relate to the fact that E2FB is more tightly associated to DREAM complexes than E2FA (Magyar et al., 2016; Lang et al., 2021). The function of these complexes is to bring together transcription factors that control G1/S genes (E2Fs) with the transcription factors controlling G2/M genes (MYB3Rs), which is essential for the timely succession of transcriptional waves during the cell cycle and entry into quiescence during differentiation (Magyar et al., 2016). DREAM complexes could thus be required to maintain the proliferative capacity of cells during replication stress. However, *e2fa-2 e2fb-1 sog1* triple mutants were completely unable to grow in the presence of replication stress, indicating that E2FA also contributes to this cellular response, maybe because it can partially compensate for the absence of E2FB. Importantly, the *e2fa-2* allele has been described as missing the transactivation and RBR1-interaction domain but retaining the “marked box” domain, which in mammals can provide a secondary interaction interface with RBR1 (Horvath et al., 2017), and still represses expression of DDR genes such as *BRCA1*, in contrast to the *e2fa-1* allele that misses this interaction domain. We therefore cannot rule out the possibility that the dramatic phenotype of the *e2fa-2 e2fb-1sog1* mutant could be due to the inhibition of E2F target genes through binding of the residual E2FA in a complex with RBR1.

### Complex transcriptional networks underlie the role of E2FA/B and SOG1 in the replication stress response

A remarkable finding in this study is that activating E2FA/B and SOG1 induces a large set of common targets. Dual control of DDR genes by activating E2FA/B and SOG1 may allow fine-tuning of the gene expression level according to the replication stress intensity. E2FA/B activity might account for basal induction levels during the S-phase, when cells are expected to be most sensitive to replication inhibitory stresses, whereas SOG1 might account for further activation in response to fork stalling. Likewise, in the *e2fa-2 e2fb-1* double mutant, we did not only see many E2F target genes being transcriptionally repressed, but an even higher number of target genes to be induced, consistent with the notion that E2FA and E2FB largely function as corepressors by recruiting RBR1 to their targets (Gombos et al., 2022). 43 of these genes are SOG1 target genes, including *BRCA1* and *RAD51*. We hypothesize that these E2F target genes are essential for S-phase progression as well as for repair of stalled replication forks, and that in the *e2fa-2 e2fb-1* plants, SOG1 is activated because of replication defects. Such dual control of target genes by both E2FA/B and SOG1 might explain the additive effects of the *e2fb-1* and *sog1* mutations seen on growth and cell cycle progression in the *pol2a* mutant, as well as the increased sensitivity of the *e2fb-1 sog1* and *e2fb-2 sog1* double and *e2fa-2 e2fb-1 sog1* triple mutants towards HU. It probably also accounts for the almost complete stalling of cell cycle progression of the *e2fa-2 e2fb-1 sog1* triple mutant in the presence of HU. Another hypothesis that could explain the hypersensitivity of the *e2fa-2 e2fb-1 sog1* triple mutant to HU is the role of E2FA in DSB repair (Biedermann et al., 2017; Horvath et al., 2017). Defects in the replication stress response triggered by SOG1 and E2FB deficiency could lead to a failure of fork stabilization mechanisms and accumulation of DSBs, which would require E2FA activation for repair.

Thus, the transcriptional network activated under replication stress is likely to be quite complex. Indeed, although E2FB and SOG1 share target genes, they seem to have opposite effects on the expression of a significant proportion of their common targets. According to our transcriptomic analysis, we can distinguish three classes of genes amongst the SOG1-regulated DDR genes (Figure 9B). A first set of genes seems to depend almost exclusively on SOG1 for their expression, as their induction is lost in *pol2a sog1* end *e2fb-1 pol2a sog1* plants (Figure 9B, class A genes). Consistently, this group of genes (corresponding to cluster 1 on Figure 5), is the least enriched in E2FA/B targets. By contrast, class B genes appear to be antagonistically regulated by SOG1 and E2FB, since their expression is lower in *pol2a sog1* mutants than in *e2fb-1 pol2a sog1* mutants. This is the case for example for *ANAC044* and *ANAC085* (Figure 9B, inset). Together with the fact that these two genes are also identified as direct RBR1 targets, this observation suggests that the repressor role of the E2FB–RBR1 complex, which we have recently described to be essential for the maintenance of cell cycle quiescence (Gombos et al., 2022), also plays an important role in the fine-tuning of the plant DDR. Finally, a large set of DDR and cell-cycle related genes, among which *WEE1*, were upregulated in the *pol2a* background, even in the absence of both E2FB and SOG1 (class C genes), although SOG1 was required for the full induction of their expression, suggesting the involvement of a third partner, likely E2FA (Figure 9B).

This observation is reminiscent of the function of E2Fs during the replication stress response in mammalian cells. Indeed, in the absence of replication stress, a negative feedback loop between the repressor E2F6, which accumulates in late S, and activating E2Fs, promotes the expression of E2F targets involved in DNA synthesis such as PCNA (Pennycook et al., 2020). In response to replication stress, the checkpoint kinase Chk1 phosphorylates and inhibits E2F6 (Bertoli et al., 2013), which allows activating E2Fs to promote the expression of major replication, repair and checkpoint effectors (Bertoli et al., 2016). This mechanism likely avoids an excessive delay in S-phase progression and accumulation of DNA damage due to fork collapse. Although our observations point to a critical role of E2FB in the control of the G2/M transition after replication stress, we cannot rule out that it could also be required to allow S-phase progression despite replication stress. Such a hypothesis would match the observation that the increase in the proportion of EdU-labeled cells during cumulative EdU experiments was extremely slow in *e2fb-1 pol2a sog1* triple mutants. Thus, besides its likely role in the control of the G2/M transition, E2FB could also function as a positive regulator of fork progression, and its loss of function might aggravate the replication defects of *pol2a* mutants.

### Emerging roles of E2FA and E2FB in the plant DNA damage response

Together, our results point to a unique role of E2FB in the plant cell’s response to replication stress. Interestingly, there is accumulating evidence that plant E2Fs are involved in the maintenance of genome integrity and play essential roles in several aspects of the DDR and even DNA repair, consistent with the functions of their animal counterpart. Indeed, both E2FA (Lang et al., 2021) and RBR1 form foci at DSBs and function independently of SOG1 to promote their repair, likely through their ability to interact with DNA repair proteins (Biedermann et al., 2017; Horvath et al., 2017). In addition, genome-wide identification of target genes revealed that RBR1 controls a large set of DDR genes (Bouyer et al., 2018), suggesting that E2F–RBR1 complexes may both control the expression of DDR genes and directly contribute to DNA repair. The respective roles of E2FA/B in the cellular response to DSBs are beginning to be unraveled, and both factors seem to contribute, E2FA by promoting DNA repair (Biedermann et al., 2017; Horvath et al., 2017), and E2FB possibly by triggering cell cycle arrest, although the two *e2fb-1* alleles do not affect this process in the same way (Lang et al., 2021). Conversely, our results suggest that E2FB is more prominently involved in the cellular response to replication stress in parallel to SOG1, and that E2FA plays only a minor role, possibly because E2FB can substitute for its activity. Only in the absence of E2FB did the *e2fa* mutation trigger sensitivity to replication stress, correlated with a longer S-phase and cell cycle duration that depended on SOG1 activity. Recently, E2FA and E2FB were shown to play distinct roles in the UV-B response (Gómez et al., 2022), as previously shown for E2FC (Gómez et al., 2019). It is therefore likely that plants E2Fs are involved in many aspects of the DDR to promote genome integrity and avoid complete cell cycle arrest triggered by DNA stress. Our transcriptome analysis reveals the extreme complexity of the transcriptional networks involving E2Fs in response to replication stress. We have only scratched the surface of the process, but more refined studies in the future will be required to understand the sequence of events occurring at the gene expression level, and how the interplay between E2FA, E2FB, SOG1 and potentially other E2F family members allows the exquisite regulation of cell cycle and DNA repair genes to maintain growth without compromising genomic integrity.

## Material and methods

### Plant material and growth conditions

All *Arabidopsis thaliana* mutant lines used in this study are in the Columbia-0 (Col-0) background and have been described previously: *e2fa-1* (*MPIZ_244*), *e2fa-2 (GABI-348E09), e2fb-1 (SALK_103138)* and *e2fb-2 (SALK_120959)* mutants were first described in Berckmans et al. (2011a, 2011b). Except for analysis of *WEE1* promoter activity, the *sog1-1* mutant was isolated in the L*er* background (Yoshiyama et al., 2009) but later introgressed in the Col-0 background and was a kind gift of Anne Britt. The *sog1-101* allele was described in Ogita et al. (2018). The *pol2a-4* mutant has been described in Yin et al. (2009) and further characterized in Pedroza-Garcia et al. (2017).

Seed were sterilized using 5 mL bayrochlore™ and 45 μL of absolute ethanol for 7 min, then washed three times with sterile water and kept at 4°C for 2 days. Seeds were sown on commercially available 0.5× Murashige and Skoog (MS, Duchefa) medium solidified with 0.8% agar (Phyto-Agar HP696, Kalys). Then, plates were transferred to long days (16-h light, 8-h dark, 21°C) in an *in-vitro* growth chamber. After 2 weeks, plantlets were transferred to soil, kept in short day conditions (8-h light at 20°C, 16-h dark at 18°C) for a week and then transferred to a long-day growth chamber (16-h light, 8-h dark, 21°C).

### Generation of reporter lines

To construct the transcriptional reporter *pWEE1-GUS*, the full-length promoter region of the *WEE1* gene was PCR-amplified (593 bp upstream of the translational start codon) and cloned into the pDONRP4-P1R entry vector by Gateway BP reaction. Site-directed mutagenesis was carried out to mutate the E2F-binding site GCGCGCAA at the −75 bp position to AACACTGT. Subsequently, both the WT (*pWEE1-FL*) and mutated (*pWEE1-mE2F*) promoter were transferred into the pMK7S*NFm14GW,0 destination vector by Gateway LR reaction. Both constructs were transferred into the *Agrobacterium tumefaciens* C58C1RifR strain harboring the pMP90 plasmid. The obtained Agrobacterium strains were used to generate stably transformed Arabidopsis lines with the floral dip transformation method (Clough and Bent, 1998).

### Root growth assay

Seeds were germinated on 0.5x MS medium and after 4 days, seedlings were transferred to fresh plates of 0.5x MS medium or 0.5x MS supplemented with 1 mM hydroxyurea (HU).

Plates were kept in a vertical position for about 2 weeks under long-day conditions. After 2 weeks, plates were scanned and images were used to measured root length by Fiji software (https://imagej.net/Fiji).

### Flow cytometry

Flow cytometry was done on flower buds of *e2f* combination mutants. Flowers buds were chopped with the help of a razor blade. Then 1 mL of nuclei isolation buffer (45 mM MgCl_2_, 30 mM sodium citrate, 60 mM MOPS, 1% (w/v) polyvinylpyrrolidone 10,000 pH 7.2), containing 0.1% (w/v) Triton X-100, and supplemented with 5 mM sodium metabisulphite and RNAse (5 U/mL). The solution was filtered and propidium iodide was added to the solution to a final concentration of 50 μg/mL. The DNA content of 5,000 to 10,000 stained nuclei was determined using a Cyflow SL3 flow cytometer (Partec-Sysmex) with a 532-nm solid-state laser (30 mW) excitation and an emission collected after a 590-nm long-pass filter. For cell cycle analysis, we used the algorithm available in the FloMax software (flomax.software.informer.com).

### EdU labeling

Seeds were germinated on 0.5x MS medium and five-day-old seedlings were transferred to 0.5x MS medium supplemented with 10 μM EdU. Plantlets were fixed at different timepoints (3, 5, 6, 9 and 11 h) with 4% (w/v) paraformaldehyde (PFA) dissolved in PME buffer (50 mM PIPES pH 6.9, 5 mM MgSO_4_, 1 mM EGTA) for 15 min under vacuum. After that, plantlets were washed twice with PME to remove the traces of PFA. Squares were drawn on polysine slides using a hydrophobic marker, root tips were cut in a drop of PME under a stereomicroscope. The PME solution was then replaced by an enzyme solution (1% (w/v) cellulase, 0.5% (w/v) cytohelicase, 1% (w/v) pectolyase in PME), and samples were incubated for 1 h in a humid chamber at 37°C. Root tips were then washed three times with 1x PME. After removing most of the liquid, root tips were squashed under a coverslip. Slides were immersed in liquid nitrogen for about 15 s, after which the coverslip was carefully removed. Slides were dried overnight. The next day, slides were washed with 1x PBS (Phosphate Buffer Saline, Sigma) and then with 3% BSA (w/v) prepared in 1x PBS. Samples were permeabilized in 0.5% Triton dissolved in 1x PBS for 30 min. Slides were washed twice with 3% BSA+1x PBS, and then the samples were incubated with Click-iT Plus EdU Alexa Fluor 488 Imaging Kit (Thermo Fisher Scientific) according to the manufacturer’s instructions, for 30 min in the dark at room temperature. Once washed with PBS 1x (pH 7.4) + BSA 3%, the nuclei were stained with Hoechst 33342 (1 μg/mL). Slides were mounted in Vectashield® and observed using an epifluorescence microscope equipped with an Apotome module (AxioImager Z.2; Carl Zeiss) fitted with a metal halide lamp and the appropriate filter sets for imaging DAPI and Alexa 488 dyes. Images were acquired with a cooled CCD camera (AxioCam 506 monochrome; Carl Zeiss) operated using the Zen Blue software (Carl Zeiss).

### Meristem length measurement

For measuring the apical root meristem, 7-day-old root tips were stained with 10 μM PI for about 5 min, and then observed with a Zeiss LSM 880 laser scanning confocal microscope using a 561-nm laser for excitation. Fluorescence was acquired between 470 nm and 700 nm. Representative images were collected from 10 to 15 roots with three biological replicates.

### GUS staining and leaf microscopy

For GUS staining, whole seedlings were stained in a 6-well plate (Falcon 3046; Becton Dickinson) as described (Beeckman and Engler, 1994). Briefly, plants were fixed in an ice-cold 80% (v/v) acetone solution for 30 min. Samples were washed three times with phosphate buffer (14 mM NaH_2_PO_4_ and 36 mM Na_2_HPO_4_) before being incubated in staining buffer (0.5 mg/mL 5-bromo-4-chloro-3-indolyl-b-D-glucuronic acid, 0.165 mg/mL potassium ferricyanide, 0.211 mg/mL potassium ferrocyanide, 0.585 mg/mL EDTA pH8, and 0.1% (v/v) Triton-X100, dissolved in phosphate buffer) at 37°C for 1 h. Samples mounted in lactic acid were observed and photographed with a stereomicroscope (Olympus BX51 microscope).

To calculate leaf parameters, the first leaves of 21-day-old plants were harvested and incubated overnight in ethanol to remove chlorophyll. Subsequently, they were cleared in lactic acid and mounted on slides, and abaxial epidermal cells (∼150 cells) were drawn in three biological repeats with a binocular microscope fitted with a drawing tube and a differential interference contrast objective. Average epidermal cell number and pavement cell size were calculated as described previously (Andriankaja et al., 2012).

### RNA extraction, RNA-Seq library preparation and quantitative RT-PCR

For RNA-Seq on shoot apices, total RNAs were extracted from the shoot apex (first 2 leaves and meristematic zone) or 30 seven-day-old plantlets, using the RNA-Plus kit (Macherey-Nagel), and libraries were prepared with 1μg of total RNA using the NEBNext® Ultra™ II RNA Library Prep Kit for Illumina® according to the manufacturer’s instructions. For RNA-Seq experiments performed on root tips, the first 2 mm of 7-day-old seedlings were collected in liquid nitrogen. RNA from samples was extracted using the RNeasy Plant Mini kit (QIAGEN) and cDNA was prepared from 1 μg of RNA using the iScript cDNA synthesis kit (Bio-Rad), both according to the manufacturer’s protocols. Libraries were sequenced on a HiSeq2000 or NextSeq500 75-bp single-end run.

Quantitative RT-PCR was performed in a final volume of 5 μl with SYBR Green I Master (Roche) and analyzed with a Lightcycler 480 (Roche) or LC96 (Roche). For each reaction, three biological and three technical repeats were performed. Primers used in this study are listed in Table S9.

### RNA-Seq data analysis

Single-end sequencing of RNA-Seq samples were trimmed using Trimmomatic-0.38 (Bolger et al., 2014) with the parameters: Minimum length of 30 bp; Mean Phred quality score greater than 30; Leading and trailing bases removal with base quality <5. Bowtie2 aligner (Langmead and Salzberg, 2012) was used for mapping to TAIR11 genome assembly. Raw read counts were used to identify differentially expressed genes using the DiCoExpress package (Lambert et al., 2020).

### Chromatin immunoprecipitation followed by high-throughput sequencing (ChIP-seq) assay

ChIP-seq was done on 2-week-old plantlets expressing the E2FB-GFP fusion. Plantlets were crosslinked in 1 % (v/v) of formaldehyde for 15 min. Crosslinking was then quenched with 125 mM glycine for 5 min. Crosslinked plantlets were grounded in liquid nitrogen and nuclei were isolated in Nuclei Lysis Buffer (0.1% SDS, 50 mm Tris-HCl pH 8, 10 mm ethylene diamine tetra acetic acid EDTA pH 8). Chromatin was sonicated for 7 min using Covaris S220 (Peak Power: 175, cycles/burst: 200. Duty Factory: 20). The sonicated chromatin was then immuno-precipitated using anti GFP antibodies (abcam, ab290), incubated at 4°C overnight with rotation on a rotating wheel. Immunocomplexes were recovered with 40 μL of Dynabeads protein A (Invitrogen, 10002D) and incubated for 2 h at 4°C with rotation. Immunoprecitated material was washed 6 times for 5 min with ChIP dilution buffer (1.1% Triton X-100, 1.2 mM EDTA, 16.7 mM Tris-HCl pH8, 167 mM NaCl, protease inhibitors) and twice in TE (1 mm Tris-HCl pH 8, 1 mM EDTA pH 8). ChIPed DNA was eluted by two 15-min incubations at 65°C with 200 μL of freshly prepared elution buffer (1% SDS, 0.1 m NaHCO_3_). Chromatin was reverse crosslinked by adding 16 μL of 5 M NaCl, incubated overnight at 65°C. The next day, chromatin was treated with RNase and Proteinase K, incubated for 3 h at 50°C, and DNA was extracted with phenol-chloroform. Ethanol was used to precipitate DNA in the presence of glycoblue and was then resuspended in 10 μL of nuclease free water. Libraries were then generated using 10 ng of DNA with NEBNext Ultra II DNA Library Prep Kit for Illumina (NEB). The quality of the libraries was assessed with Agilent 2100 Bioanalyzer Instrument, and the libraries were subjected to 1 × 7 bp high-throughput sequencing by NextSeq 500 (Illumina).

### Statistical analysis

Statistical analyses were performed as indicated in the figure legends.

### Accession numbers

RNA-Seq raw data from this study were deposited in Gene Expression Omnibus (accession number GSE220849 for RNAseq data obtained on root tips of e2fa-2 e2fb-1 mutants, and GSE220872 for transcriptome data obtained in *pol2a, e2fb-1 pol2a* and *e2fb-1 pol2a sog1* mutants). Sequence data from this article can be found in the Arabidopsis Genome Initiative or GenBank/EMBL databases under the following accession numbers: *E2FA* (At2g36010); *E2FB* (At5g22220); *POL2A* (AT1G08260); *SOG1*(AT1g25580).

## Supplemental data

The following materials are available in the online version of this article.

**Supplemental Table S1: List of E2FA (S1A), E2FB (S1B) targets and their shared targets with SOG1 (S1C)**.

**Supplemental Table S2: DEG in *pol2a, pol2a sog1* and *e2fb-1-1 pol2a sog1* mutants**

**Supplemental Table S3: Overlaps between up-regulated genes in the analyzed mutants**

**Supplemental Table S4: List of genes found in each cluster of Figure 5C (Table S4A) and proportion of E2F targets in each cluster (Table S4B)**

**Supplemental Table S5: List of DEG identified in the RNA-seq analysis performed on root tips of *e2fa-2 e2fb-1* double mutants: upregulated genes (Table S5A), downregulated genes (Table S5B) and list of upregulated genes targeted by E2FA and/or B (Table S5C)**.

**Supplemental Table S6: List of DNA-replication-related genes down-regulated in *e2fa-2 e2fb-1* double mutants**

**Supplemental Table S7: GO analysis of down-regulated genes in *e2fa-2 e2fb-1* double mutants**

**Supplemental Table S8: GO analysis of up-regulated genes in *e2fa-2 e2fb-1* double mutants**

## Acknowledgments

The authors thank Etienne Delannoy at Marie-Laure Martin-Magniette (IPS2) for advice on RNA-Seq data analysis; and Annick Bleys for critical reading and editing of the manuscript. Imaging experiments conducted at IPS2 benefited from the Imaging Facility of the Institute.

## FIGURE LEGENDS

**Supplemental Figure S1.**
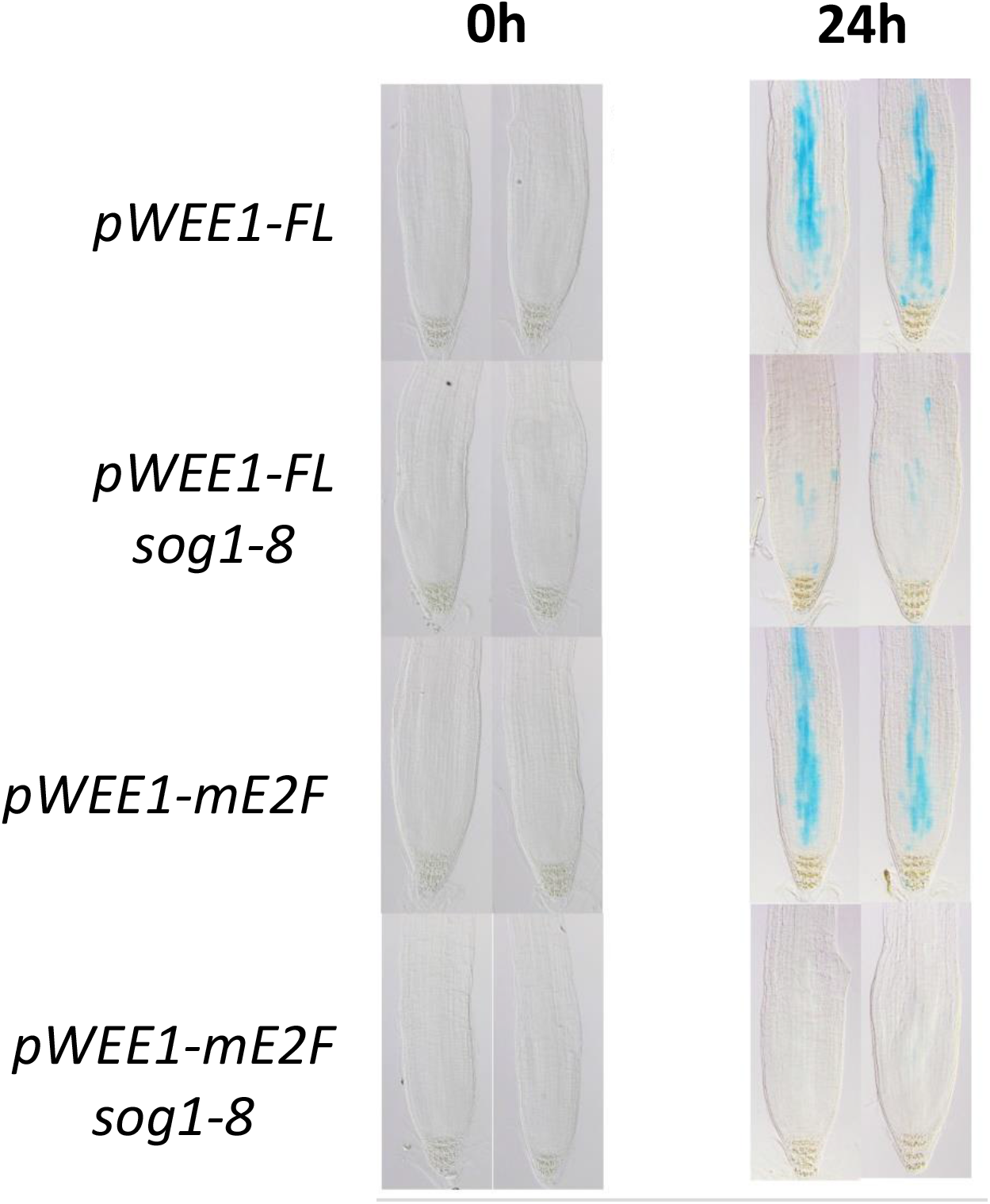
*WEE1* activation in response to hydroxyurea relies both on SOG1 and E2Fs. Constructs encompassing the full-length (FL) *WEE1* promoter driving the expression of the beta-glucuronidase (*GUS*) gene were introduced into wild-type or *sog1-101* mutants. GUS staining was observed after 24 h of treatment with the replicative stress-inducing drug hydroxyurea (1 mM). Staining was drastically reduced but still visible in the *sog1-8* background. However, when the E2F binding site was deleted (mE2F), residual activation was lost, demonstrating that E2Fs can contribute to WEE1 activation in response to replication stress.

**Supplemental Figure S2.**
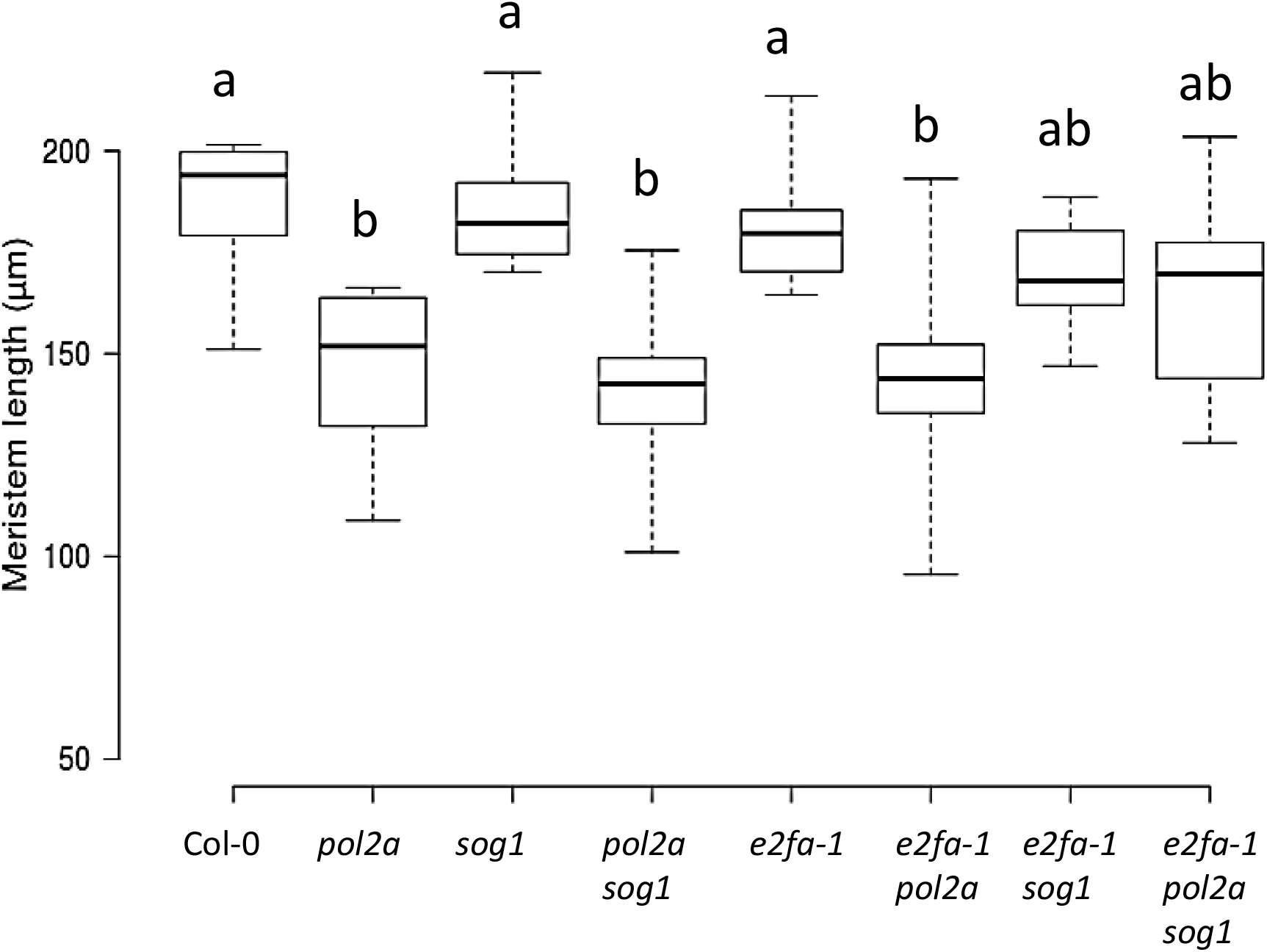
E2FA does not affect apical root meristem length in the *pol2a* background. Data is taken from at least ten roots and are representative of two independent experiments. Different letters indicate statistically significant differences (ANOVA and Tukey test p < 0.01).

**Supplemental Figure S3.**
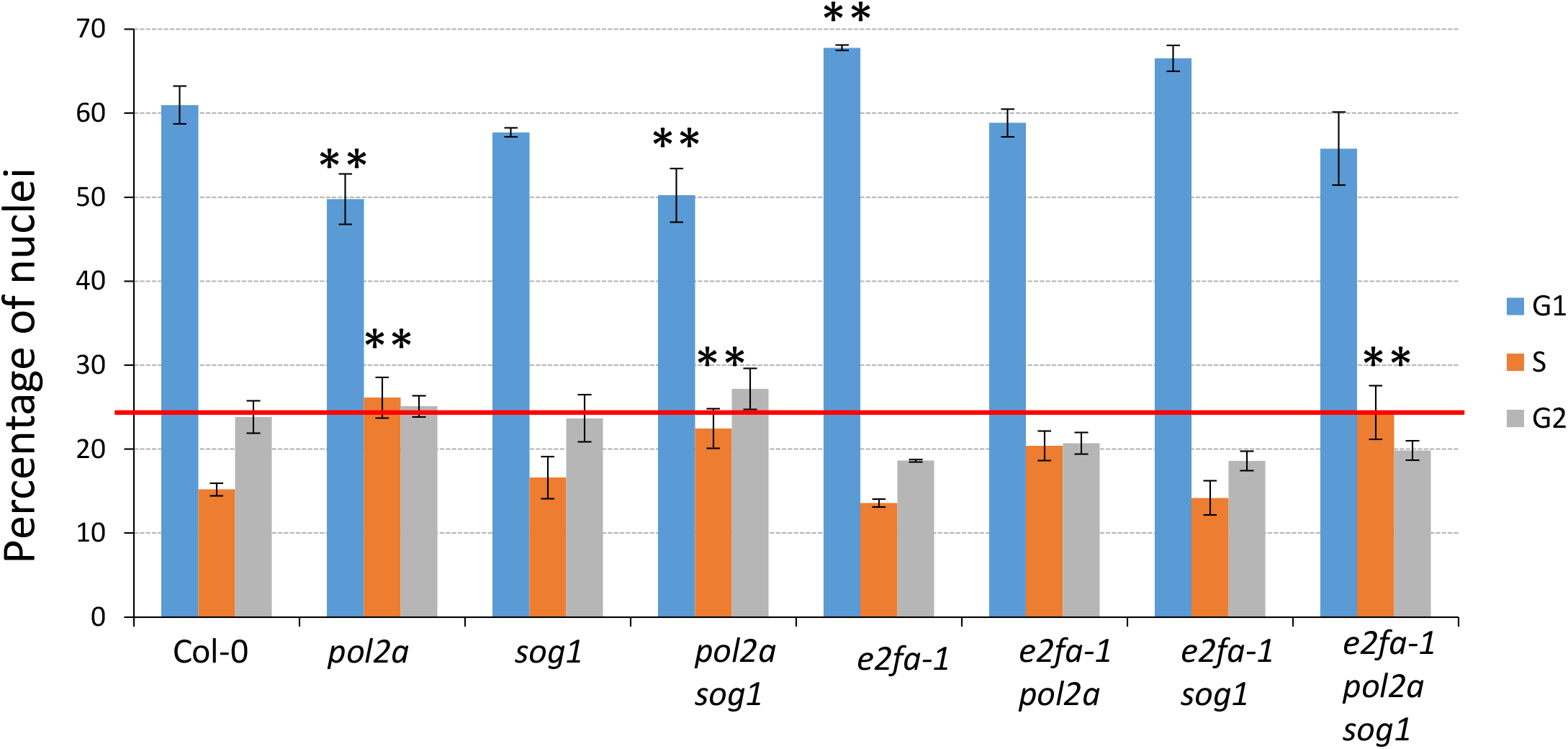
Loss of E2FA does not affect cell cycle progression in *pol2a* or *pol2a sog1* mutants. Flow cytometry was used to analyze the cell cycle phases in the flower buds of Col-0, *pol2a, sog1, pol2a sog1, e2fa, e2fa pol2a, e2fa sog1 and e2fa pol2a sog1* mutant lines. For each cell cycle phase, the percentage of nuclei was calculated. Values are average ± SD. Asterisks denote significant differences with respect to percentages observed in the wild type (Kruskal-Wallis test, * p < 0.05, ** p < 0.01). Data are representative of three biological replicates. The red line indicates the proportion of G2 nuclei in the wild-type.

**Supplemental Figure S4.**
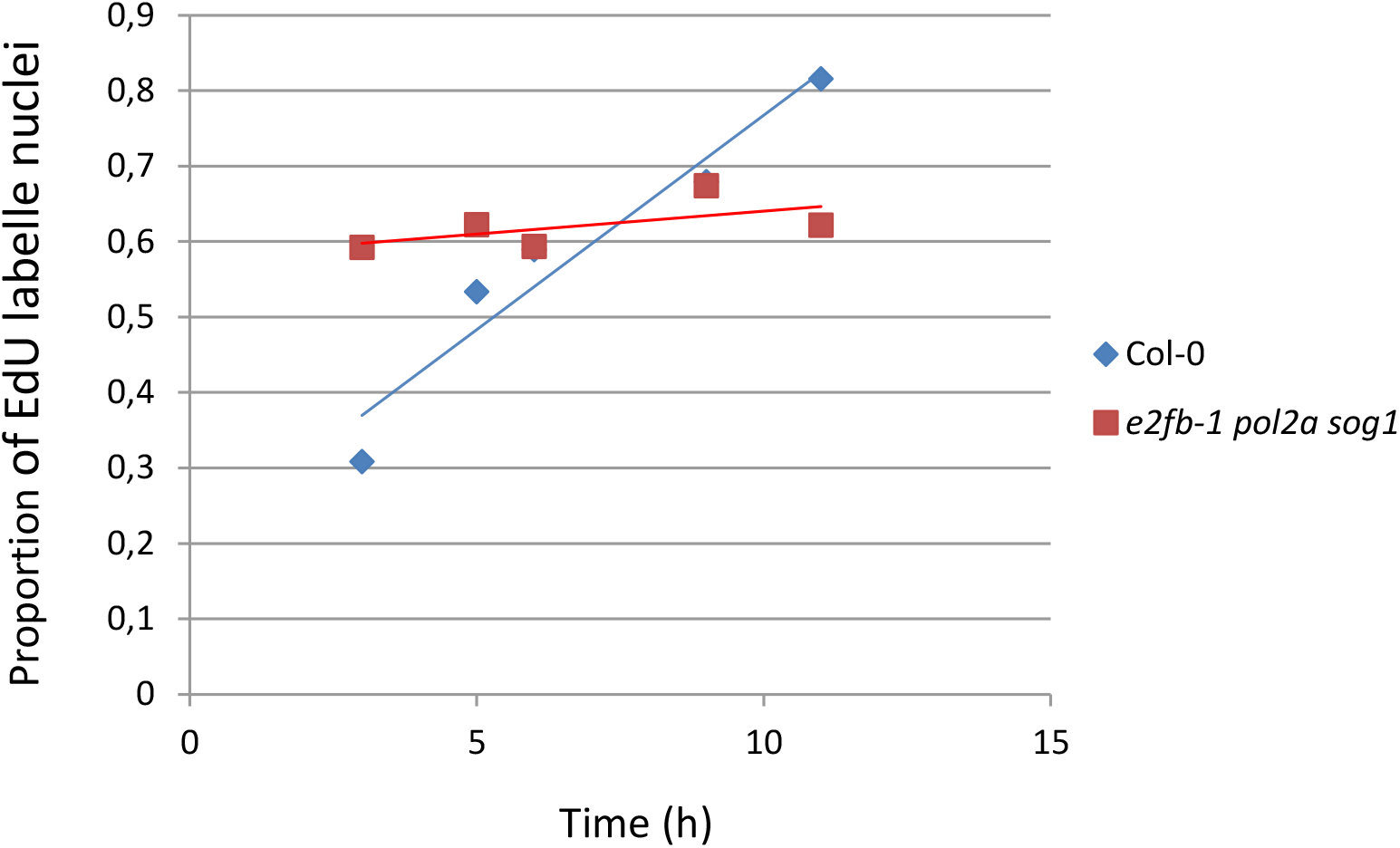
Cumulative EdU incorporation suggests that cell cycle progression is extremely slow in *e2fb-1 pol2a sog1* mutants. Cumulative EdU incorporation was performed on wild type (Col-0) and *e2fb-1 pol2a sog1* triple mutants for 12 h. At each timepoint, the proportion of EdU-positive nuclei was estimated through microscopic analysis (n > 1000). The proportion of EdU-labeled nuclei was very high at early timepoints in the triple mutant, indicating that the S-phase duration represents a high proportion of the total cell cycle length. Furthermore, this proportion remained stable throughout the kinetics, indicating that progression through the cell cycle is very slow.

**Supplemental Figure S5.**
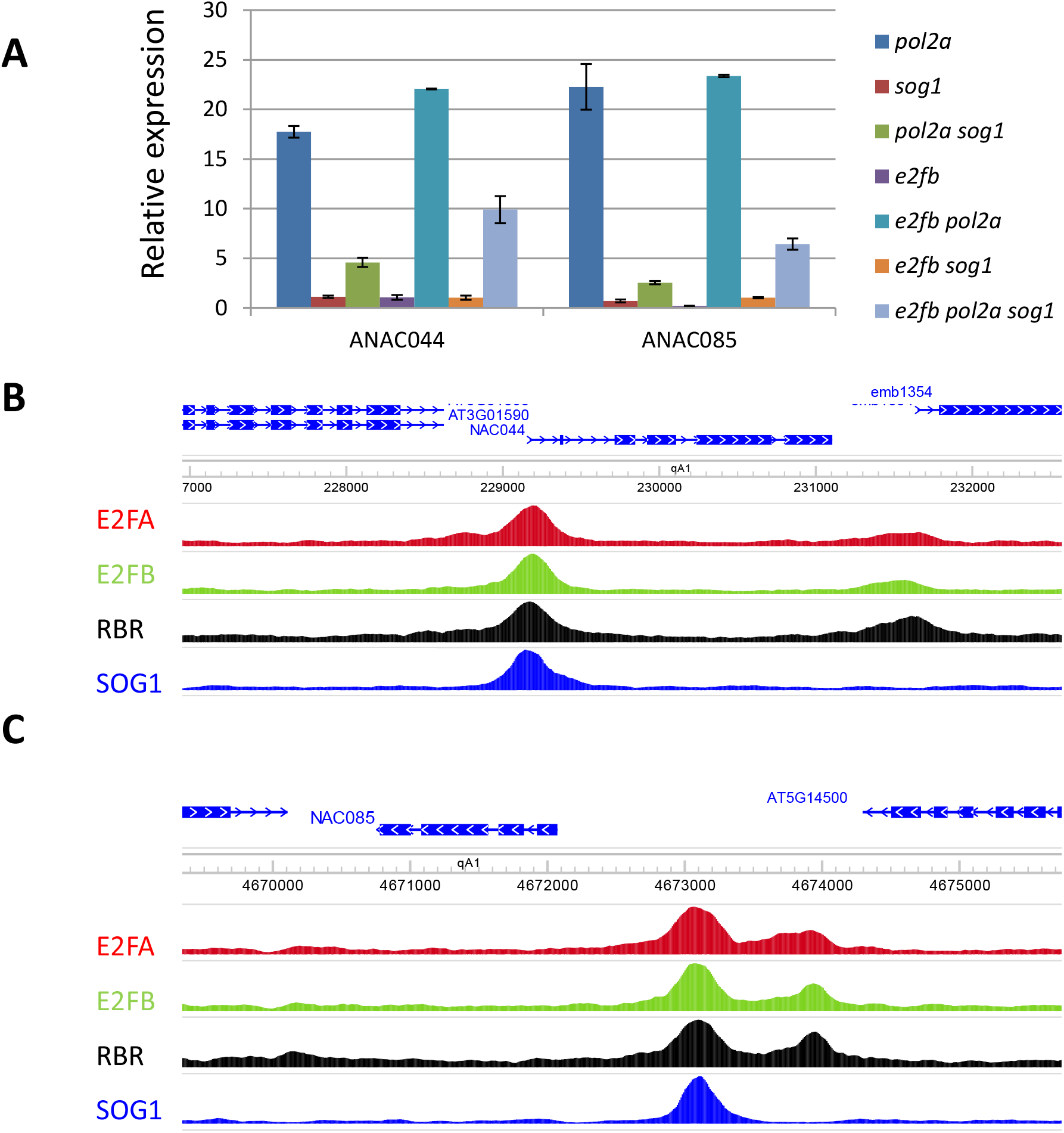
Expression of *ANAC044* and *ANAC085* is under the control of SOG1 and E2FA/B. A: RT-qPCR analysis of *ANAC044* and *ANAC085* expression in the indicated mutant lines. RNA was extracted from shoot apices of 5-day-old plantlets. Expression was normalized with the *TIP4* and *SAM* genes as described in Gentric et al. (2020). B: Screenshot of ChIP-seq data showing E2FA, E2FB, RBR and SOG1 binding on the promoter of *ANAC044*. C: Screenshot of ChIP-seq data showing E2FA, E2FB, RBR and SOG1 binding on the promoter of *ANAC085*.

**Supplemental Figure S6.**
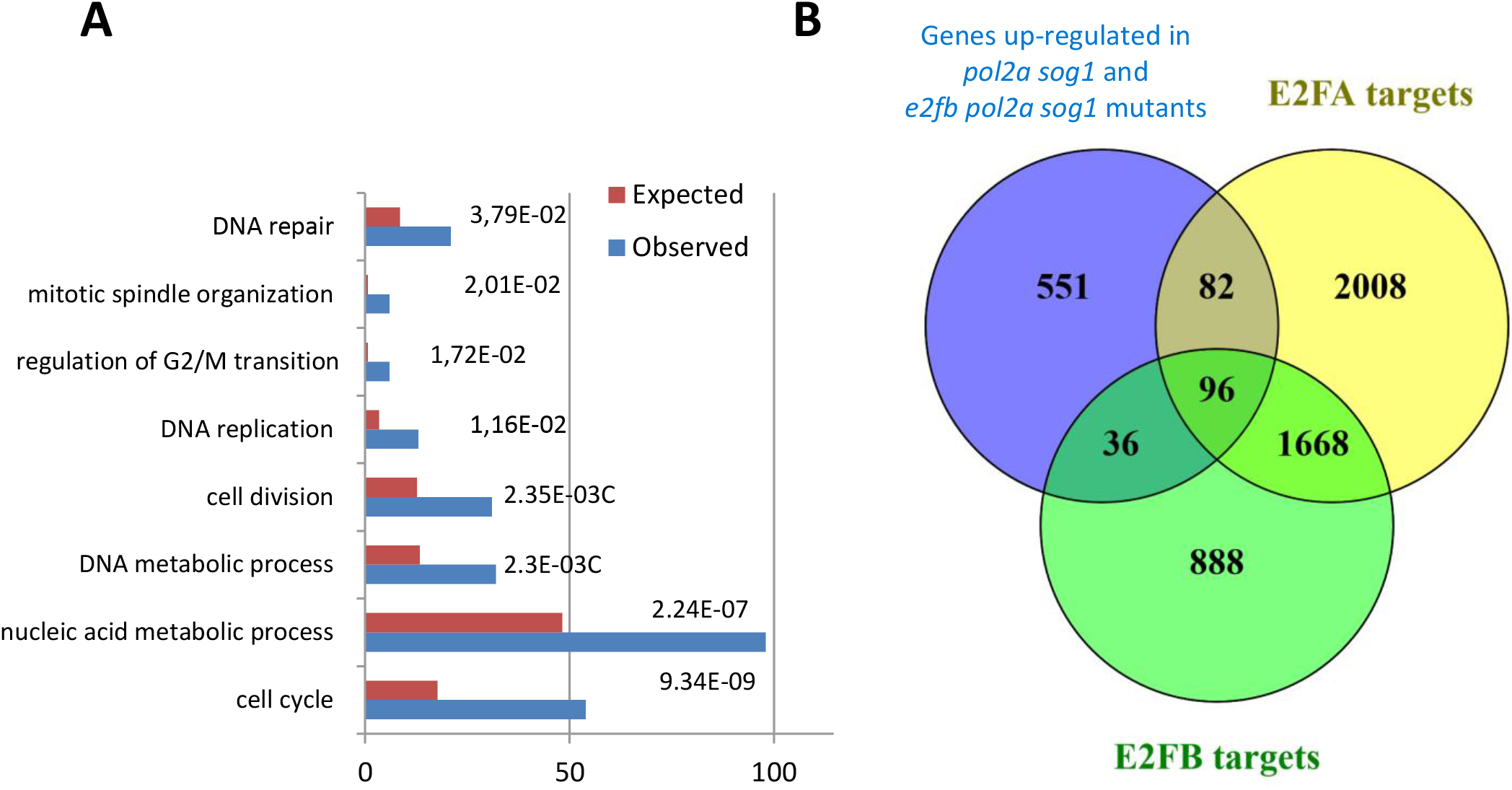
Genes specifically upregulated in *pol2a sog1* and *e2fb-1 pol2a sog1* are enriched in cell cycle-related genes and E2Fa targets. A: GO analysis of the 765 genes upregulated in *pol2a sog1* and *e2fb pol2a sog1* but not in *pol2a* mutants. B: Overlap between this list of genes and the list of E2FA and E2FB target genes.

**Supplemental Figure S7.**
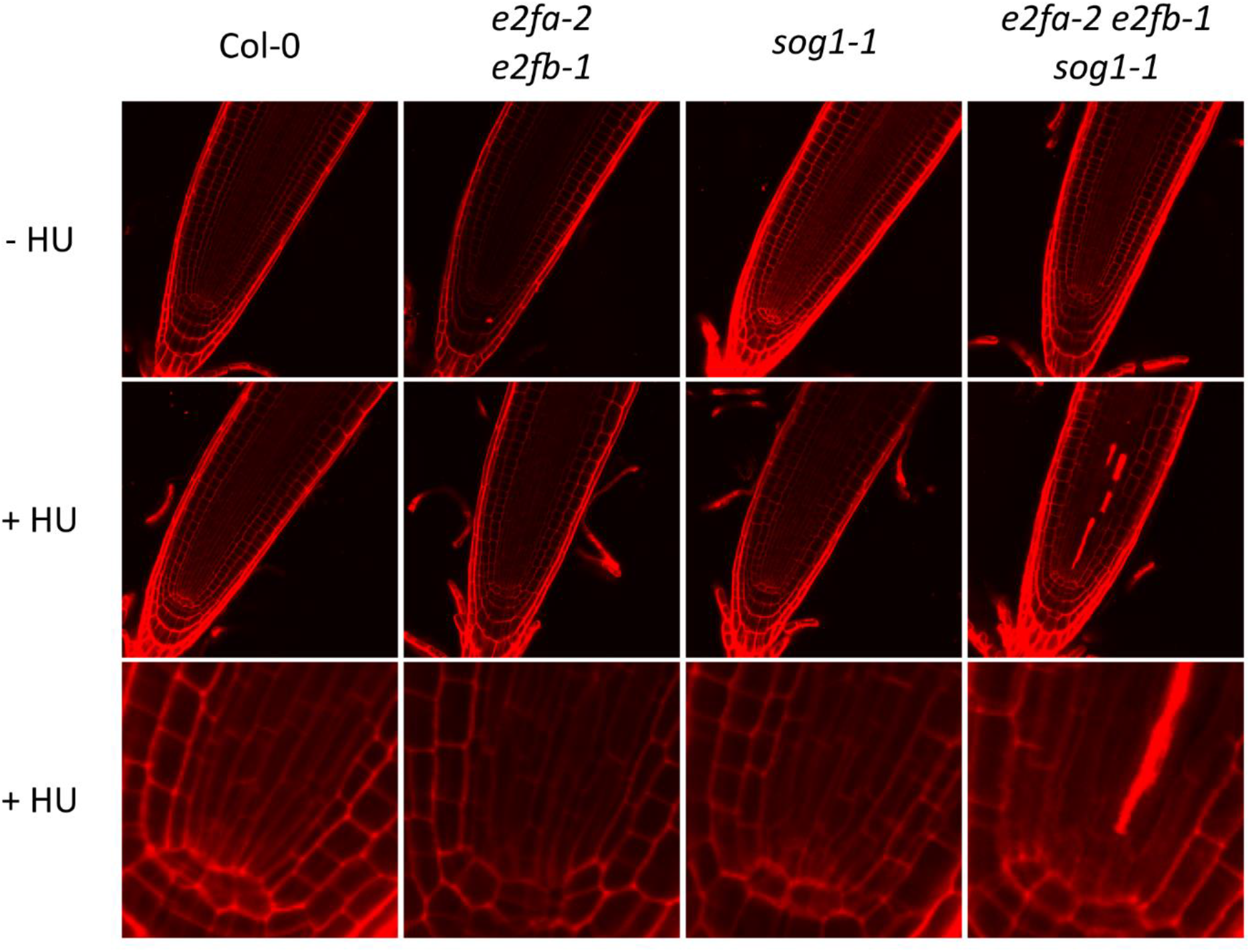
*e2fa-2 e2fb-1 sog1* triple mutants are hypersensitive to replication stress. Confocal images of 7-day-old *e2fa-2 e2fb-1 sog1* seedlings transferred for 24 h to control medium (-HU) or medium containing 1 mM HU (+ HU), stained with propidium iodide. The bottom row is a close-up of stem cell area of the middle row.

**Supplemental Figure S8.**
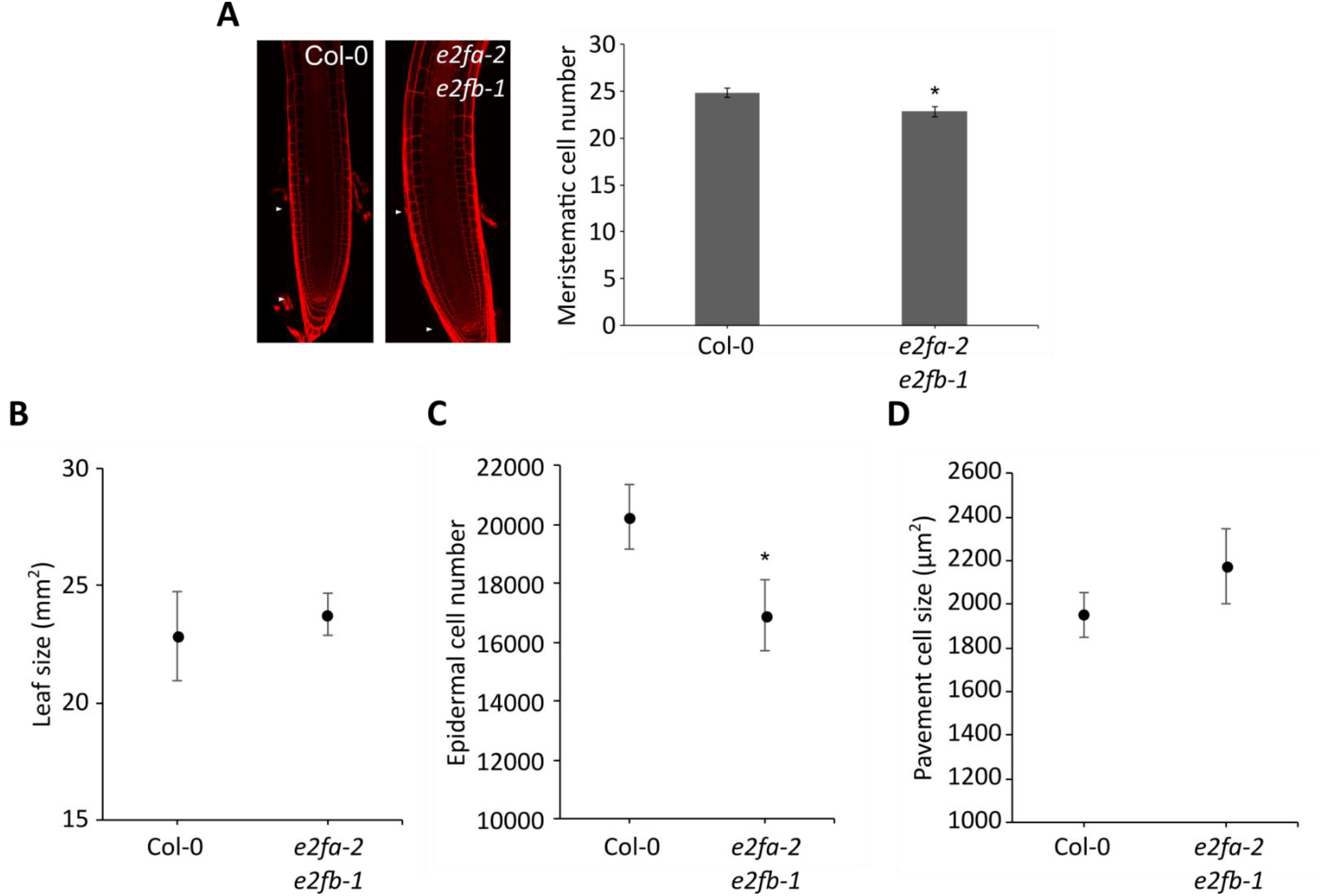
The *e2fa-2 e2fb-1* double knockout tends to have fewer cells in both root meristem and adult leaves. A: Left: Confocal images of the root tips of 7-day-old wild-type (Col-0) and *e2fa-2 e2fb-1* seedlings. Right: Meristematic cortical cells were counted in the roots. Data represent mean ± SEM (n > 20). B-D: Leaf size, epidermal cell number and pavement cell size were measured of leaf 1 and 2 of 21-day-old plants (n = 5). Data represent mean ± 95% confidence interval. A-D: Asterisks indicate statistical significance compared with Col-0 based on Student’s t-test; *, p < 0.05.

**Supplemental Table S1: List of E2FA, E2FB and shared SOG1 target genes**.

**Supplemental Table S1A: List of E2FA target genes investigated in this study**. This list shows target genes identified by TChAP in Verkest et al. (2014) (indicated in red) and by classical ChIP in Bourbousse et al. (2018) (enrichment > 2.4, FDR < 0.05). In both cases, peaks found in the coding sequence or further than 1 kb away from the closest transcription start site were left out.

**Supplemental Table S1B: List of E2FB target genes investigated in this study (enrichment > 2, FDR < 0.05 as described in Gombos et al. (2022))**. Peaks found in the coding sequence or further than 1 kb away from the closest transcription start site were left out.

**Supplemental Table S1C: Target genes shared between E2FA, E2FB and SOG1**. Shared E2FA and SOG1 target genes were defined as the union between E2FA targets identified in both TChAP replicates in Verkest et al. (2014) and SOG1 targets identified by ChIP-seq in Bourbousse et al. (2018) (enrichment > 2.5, FDR < 0.05). Shared E2FB and SOG1 target genes were defined as the union between E2FB targets identified by ChIP-seq in Gombos et al. (2022) (enrichment > 2.5, FDR < 0.05) and the same SOG1 targets.

**Table S2: Differentially expressed genes in *pol2a, pol2a sog1* and *e2fb-1 pol2a sog1* mutants**.

**Supplemental Table S2A: Differentially expressed genes between *pol2a sog1* mutants and the wild type (Col0)**. Genes highlighted in gray have an absolute fold change lower than 1.5.

**Supplemental Table S2B: Differentially expressed genes between *e2fb-1 pol2a sog1* mutants and the wild type (Col0)**. Genes highlighted in gray have an absolute fold change lower than 1.5.

**Supplemental Table S2C: Differentially expressed genes between *pol2a* mutants and the wild type (Col0)**. Genes highlighted in gray have an absolute fold change lower than 1.5.

**Supplemental Table S2D: Differentially expressed genes between *e2fb pol2a sog1* mutants and *pol2a sog1* mutants**. Genes highlighted in gray have an absolute fold change lower than 1.5.

**Supplemental Table S3: Overlaps between upregulated genes in the various mutant combinations**.

**Supplemental Table S4: List of genes found in each cluster of Figure 5C**.

**Supplemental Table S4A: Lists of genes found in each cluster shown on Figure 5C**. The log2fold change (FC) of expression in each mutant line compared with the wild type is indicated. E2FA targets are highlighted in yellow, E2FB targets in blue and common targets in green.

**Supplemental Table S4B: Number and percentage of E2F targets found in each cluster defined in Supplemental Table S4A**.

**Supplemental Table S5: List of differentially expressed genes identified in the RNA-Seq analysis performed on root tips of *e2fa-2 e2fb-1* double mutants**.

**Supplemental Table S5A: List of differentially expressed genes in *e2fa-2 e2fb-1* mutants**. Only genes with an absolute FC > 1.5 (i.e. log2FC > 0.585) were considered for further analysis. Genes with a smaller FC are highlighted in gray.

**Supplemental Table S5B: List E2F targets downregulated in *e2fa-2 e2fb-1* mutants. Supplemental Table S5C: List E2F targets upregulated in *e2fa-2 e2fb-1* mutants**.

**Supplemental Table S6: List of DNA-replication-related genes downregulated in *e2fa-2 e2fb-1* double mutants**.

**Supplemental Table S7: GO analysis of downregulated genes in *e2fa-2 e2fb-1* double mutants**.

**Supplemental Table S8: GO analysis of upregulated genes in *e2fa-2 e2fb-1* double mutants**.

**Supplemental Table S9: Sequences of primers used for RT-qPCR analyses**.

